# Modulation statistics of natural soundscapes

**DOI:** 10.1101/2025.03.04.638661

**Authors:** Nicole Miller-Viacava, Frédéric Apoux, Regis Ferriere, Nicholas R. Friedman, Timothy C. Mullet, Jérôme Sueur, Jacob Willie, Christian Lorenzi

## Abstract

Modulation statistics of “natural soundscapes” were estimated by calculating the modulation power spectrum (MPS) of a database of acoustic samples recorded in nine pristine terrestrial habitats for four moments of the day and two contrasting periods differing in precipitation level. Following Singh and Theunissen (2003), a set of statistics estimating low-pass quality, starriness, separability, asymmetry, modulation depth and 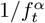 temporal-modulation power-law relationships were calculated from the MPS of the samples and related to geographical, meteorological factors and diel variations. MPS were found to be generally low-pass in shape in the modulation domain, with most of their modulation power restricted to low temporal (<10-20 Hz) and spectral modulations (<0.5-1 cycles/kHz). Modulation statistics distinguished between habitats irrespective of moment of the day and precipitation period, with a greater role of Modulation depth and Starriness. Separability and Starriness were found to be related to the global biodiversity decrease from tropical to polar regions, suggesting that the lack of joint high spectral and fast temporal modulations, and MPS complexity are important features characterising “biophony”, the collective sound produced by animals in a given habitat. These findings may help guide research on monitoring auditory behaviors and underlying mechanisms expected to exploit regularities of natural scenes.

## I. INTRODUCTION

Natural soundscapes are complex acoustic patterns produced by superimposed biological and geophysical sound sources (also referred to as “biophony” and “geophony”, respectively; Krause, 1987), shaped by the specific way sound propagates through the environment. These scenes are defined as “natural” in the sense that the contribution of human activity is expected to be minimal or negligible, as is the case for sounds recorded in protected reserves, national parks or certain rural areas (Grinfeder *et al*., 2022b; Lorenzi *et al*., 2023).

Over the last decades, a wealth of research in auditory neuroscience, neuro-ethology and psychoacoustics attempted to test fundamental principles in sensory coding derived from information theory, namely the general “correspondence principle” positing that regularities in natural sound ensembles (e.g., animal vocalisations or natural soundscapes as defined above) should be visible in the response properties of auditory neurons (Nelken *et al*., 1999) and the related efficient-coding hypothesis (e.g., He *et al*., 2023; Lesica and Grothe, 2008; Lewicki, 2002; Park *et al*., 2021; Smith and Lewicki, 2006) postulating that auditory processing has been optimized for natural sounds through development and/or evolution. These studies shared the same approach that proved to be fruitful in vision science over the last decades (Attneave, 1954; Barlow, 1961; Field, 1987, 1994; Simoncelli and Olshausen, 2001; Tolhurst *et al*., 1992; Torralba and Oliva, 2003), assuming that the auditory system of human and non-human animals –in order to process efficiently incoming sounds– should take advantage of the redundancy of natural sounds and soundscapes. Consistent with this framework, several studies showed that the rate and efficiency of information transmission of auditory neurons is highest for natural stimuli (e.g., Escabí *et al*., 2003; Garcia-Lazaro *et al*., 2006, 2011; Rieke *et al*., 1995; Woolley *et al*., 2005). This motivated the need to characterize the statistics of large natural sound ensembles such as animal vocalisations, speech sounds and so-called “environmental sounds” here-defined as sounds produced by geophysical sources (e.g., Attias and Schreiner, 1996; McDermott *et al*., 2009; McDermott and Simoncelli, 2011; McWalter and Dau, 2017; Singh and Theunissen, 2003; Theunissen and Elie, 2014). In these studies, separate acoustic recordings of animal vocalisations, forest sounds, stream, rain and fire sounds were submitted to a spectro-temporal decomposition allowing for an analysis of their spectral and temporal modulation content. Such an endeavour was supported by the demonstration of the importance of spectro-temporal modulation cues for human (e.g., Drullman, 1995; Shannon *et al*., 1995; Venezia *et al*., 2016) and non-human (e.g., Dooling and Popper, 2000; Fay and Simmons, 1999; Römer, 1998, 2001; Stebbins and Moody, 1994) animal communication, together with the demonstration of high auditory sensitivity to modulation cues in sounds for human and non-human observers (Chi *et al*., 1999; Dooling and Popper, 2000; Kohlrausch *et al*., 2000; Long, 1994; Viemeister, 1979; Wallaert *et al*., 2018) and selective tuning of cortical and subcortical neurons to spectro-temporal modulation cues relevant to communication sounds (e.g., Eggermont, 2002; Hsu *et al*., 2004; Joris *et al*., 2004; Rees and Malmierca, 2005; Schreiner and Urbas, 1986; Woolley *et al*., 2005).

Four pioneering studies conducted by Voss and Clarke (1975), Attias and Schreiner (1996) and Singh and Theunissen (2003) found that environmental sounds have most of their modulation energy restricted to slow temporal and low spectral modulations, and that animal vocalisations (e.g., bird calls) show most of spectral modulation power at relatively slow temporal modulations (< 5 Hz). These studies also showed that the average temporal components of the amplitude-modulation spectrum for environmental sounds follows a power law 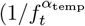 with *f*_*t*_ corresponding to the ortemporal amplitude-modulation rate) with an exponent *α*_temp_ = 1 between that of animal vocalisations (*α*_temp_ = 2) and white noise (*α*_temp_ = 0). This power-law behaviour which reveals long-term temporal correlations in the amplitude and thus, the possibility to predict portions of the acoustic signals from earlier ones, was demonstrated for a large number of species (Jermyn *et al*., 2023), for water sounds (Geffen *et al*., 2011) and soundscapes recorded in rural areas (e.g., De Coensel and Botteldooren, 2006; De Coensel *et al*., 2003). Importantly, this 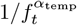 relationship implies that the second-order statistics of natural sound ensembles are “scale invariant” in the time domain, that is invariant to temporal compression or expansion, possibly contributing to the (limited) perceptual constancy reported in the temporal-modulation domain for human listeners (Ardoint *et al*., 2008; Fu *et al*., 2001; Geffen *et al*., 2011).

These pioneering studies provided invaluable information about the potential degree of statistical redundancy of natural scenes that proved crucial for advancing our understanding of auditory mechanisms and auditory perception in human and non-human animals. However, these studies used acoustic databases that were rather small in size and in most cases, their specific content was generally poorly described and documented, questioning the possibility to generalise their findings and limiting strongly in-depth analysis of the results. In addition, these databases had limited ecological validity in that they artificially over-represented certain categories of biotic and abiotic sounds (e.g., mammal sounds), and removed them from their original acoustic environment ignoring therefore the contribution of geophysical sounds (Farina *et al*., 2021) and sound propagation effects that strongly differ across closed and open habitats (e.g., forest v/s desert or savannah) (for a review of these effects see Forrest, 1994; Grinfeder *et al*., 2022b; Lorenzi *et al*., 2023). In addition, these studies did not take into account seasonal and circadian effects produced by the natural habitats that are expected to severely alter their spectro-temporal characteristics (Grinfeder *et al*., 2022b).

Large databases and availability of metadata obviously guarantee accurate estimation of “true” statistical regularities of natural scenes and interpretability of results. Ecological validity of acoustic databases plays an even greater role as it guarantees that acoustic material is representative of that experienced in real natural environments and relevant to the sensory process being investigated (Holleman *et al*., 2020). In that respect, several important features of natural soundscapes have been highlighted by previous work in soundscape ecology (e.g., Farina and Gage, 2017; Grinfeder *et al*., 2022b; Lorenzi *et al*., 2023; Pijanowski *et al*., 2011; Sueur and Farina, 2015). First, the biological component of natural soundscapes shows a strong and ubiquitous periodicity caused by the circadian cycle, with a chorus at dawn and at dusk (see Gil and Llusia, 2020), and seasonal variations (e.g., Gage and Axel, 2014). Second, annotations of large databases revealed that birds, insects and, to a lower extent, amphibians are the most predominant in natural soundscapes across a wide variety of terrestrial biomes (Chen *et al*., 2021; Diepstraten and Willie, 2021; Divyapriya and Pramod, 2019; Gasc *et al*., 2018a,b; Mullet *et al*., 2016; Phillips *et al*., 2018). Given that bird, insect and anuran sounds are quite different from mammal vocalisations with, among other, faster temporal modulations (Aubin and Bremond, 1983; Capranica *et al*., 1985; Catchpole and Slater, 2008; Fonseca, 2014; Gerhardt and Bee, 2006; Stein, 1968; Sueur and Aubin, 2003), this predominance of birds and insects imposes specific acoustic regularities to natural soundscapes, to which mammals, including human ancestors, have been exposed for at least several million years (Senter, 2008). Third, most of previous studies used isolated sounds but biological sounds are hardly heard in isolation in real-world natural settings. Within natural soundscapes, biophony is mixed up with geophony to form specific and potentially unique combinations associated with a given habitat and biome. As an example, rain may act as an acoustic masker and alter animal communication. For certain species and habitats, rain has also been found to stimulate or disrupt animal reproductive behavior. Rain therefore yields substantial changes in animal communication and biophony (Amézquita and Hödl, 2004; Bedoya *et al*., 2017; Grinfeder *et al*., 2022a; Lack, 1950; Lengagne and Slater, 2002; Tishechkin, 2013; Zina and Haddad, 2005). Finally, the biological and geophysical constituents of natural soundscapes are shaped by the specific way sounds propagate in the habitat. Because of sound scattering caused by vegetation, closed environments such as deciduous or coniferous forests tend to attenuate a specific range of audio-frequency components, creating “sound windows”, and reverberation alters strongly the transmission of fast modulations. In comparison, open environments such as grasslands, savannahs, tundras or deserts show little reverberation but alter strongly the transmission of slow modulations because of atmospheric turbulence (Forrest, 1994; Michelsen and Larsen, 1983; Morton, 1975; Richards and Wiley, 1980). The “acoustic adaptation hypothesis” coined by Morton (1975) posits that animal vocalisations are shaped by such environmental constraints with signals produced by species inhabiting the same ecosystem showing some acoustic similarity (though this hypothesis has been recently challenged, see for example Freitas *et al*., 2024). For these reasons, the importance of certain modulation statistics for animal sounds may not have been properly estimated in previous studies. The work by Mouterde *et al*. (2014) illustrates this idea by showing how the harmonic structure and fast temporal modulations of zebra-finch vocalisations are dramatically attenuated by their propagation over about 250 m in their natural environment. These features should be taken into account when searching for universal and robust statistics of natural acoustic scenes and more specifically, markers of biophony.

Thoret *et al*. (2020) presented the first attempt to characterize the temporal-modulation characteristics of soundscapes recorded in four distinct habitats of a protected nature reserve at four moments of the day and four seasons during a single year. The relatively slow spectro-temporal modulation cues were found to provide sufficient information for accurate classification of habitat (forest, grassland, chaparral and meadow), moment of the day and season using support vector machine. This was confirmed in a follow-up study conducted by Apoux *et al*. (2023), comparing discrimination capacities of convolutional neural networks to human listeners. However, both studies were limited to a single temperate biome and further work is warranted to assess the universality of these findings. The goal of the present study is to extend this initial work by characterising the modulation statistics of a larger database of natural soundscapes taking into account a greater number of ecological factors that have an impact on natural soundscapes.

To address this issue, modulation statistics of natural soundscapes were estimated and analysed by calculating the modulation power spectrum (MPS) of a large database of one-sec long acoustic samples recorded in nine different terrestrial habitats with minimal human activity, for four moments of the day and two periods of two months each differing in precipitation level. Following the approach developed by Singh and Theunissen (2003), a set of six quantifiers estimating low-pass quality, starriness, separability, asymmetry, modulation depth and temporal-modulation power relationships were calculated from the MPS of the acoustic samples and related to geographical, meteorological factors, and diel variations. These quantifiers are the modulation statistics analysed in the present work. In their seminal study, Singh and Theunissen (2003) showed clear differences between the modulation statistics of biological (speech and bird vocalisations) and geophysical (rustling brush, crunching leaves, rain, stream sounds) sound sources. Altogether, these results along with the well-known latitudinal biodiversity gradient demonstrated by ecologists – the global biodiversity decrease from tropical to polar regions (e.g., Brown *et al*., 2004; Gaston, 2000; Gaston and Blackburn, 2000; Lomolino *et al*., 2018; Rolland and Freeman, 2023)– suggest that the above modulation statistics should differ as a function of habitat, moment of the day, and precipitation period. More importantly, it is expected that these modulation statistics should distinguish natural soundscapes recorded in equatorial habitats from those recorded at other latitudes because biodiversity peaks in equatorial regions, an effect that proved to be difficult to demonstrate using current ecoacoustic indices which are not based on an explicit analysis of the spectro-temporal modulation content of sounds (Eldridge *et al*., 2018; Pan *et al*., 2024). In particular, this study aims at unveiling whether the modulation statistics extracted from the MPS can characterise and distinguish different habitats irrespective of moment of the day and precipitation period.

A characterisation of modulation statistics in this ecologically-valid context should be useful for future psychoacoustical, neuroscientific and ethological studies aiming at testing efficient-coding principles and/or identifying auditory cues and sensory mechanisms used by human and non-human animals for discriminating habitats and their global features (as in Apoux *et al*., 2023; McMullin *et al*., 2024; Miller-Viacava *et al*., 2024; Thoret *et al*., 2020). It should also be useful for ecoacoustical studies searching for robust acoustic markers of biodiversity.

## II. MATERIAL AND METHODS

### A. Natural soundscapes recordings

#### 1. Sound database

A large database of natural soundscapes was created thanks to the collaboration of nine research teams in ecoacoustics. These teams had already recorded natural soundscapes from nine different terrestrial habitats and agreed to share their recordings (De Baudouin *et al*., 2024; Gage and Axel, 2014; Grinfeder *et al*., 2022a; Mullet *et al*., 2024; Phillips *et al*., 2018; Ross *et al*., 2018). These nine habitats are spread across different continents and latitudes as shown in figure 1 where a picture and location (based on GPS coordinates) from each recording location can be seen. Natural soundscapes were recorded for long periods in natural parks or reserves where terrestrial human activity is supposed to be marginal. All the relevant information regarding recordings is detailed in table I with habitats sorted by distance to the Equator (|latitude| of each location in degrees). Amongst these nine habitats, two are open (habitats 4 and 7) and seven are closed (habitats 1, 2, 3, 5, 6, 8, and 9), with seven in the northern (habitats 1, 2, 3, 4, 5, 8, and 9) and two in the southern hemisphere (habitats 6 and 7).

**TABLE I.**
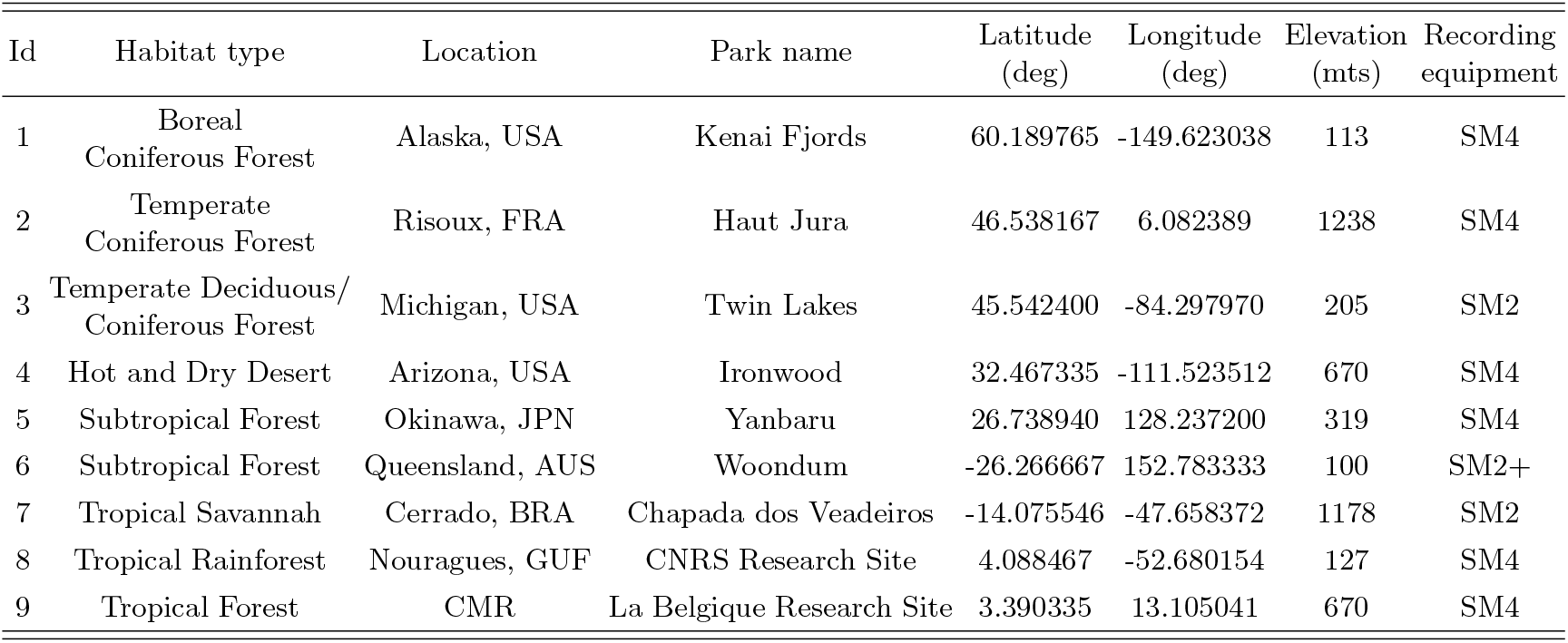
General characteristics and localisation of the nine habitats of the Hearbiodiv database. Information about recording material is shown in the last column.

**FIG. 1.**
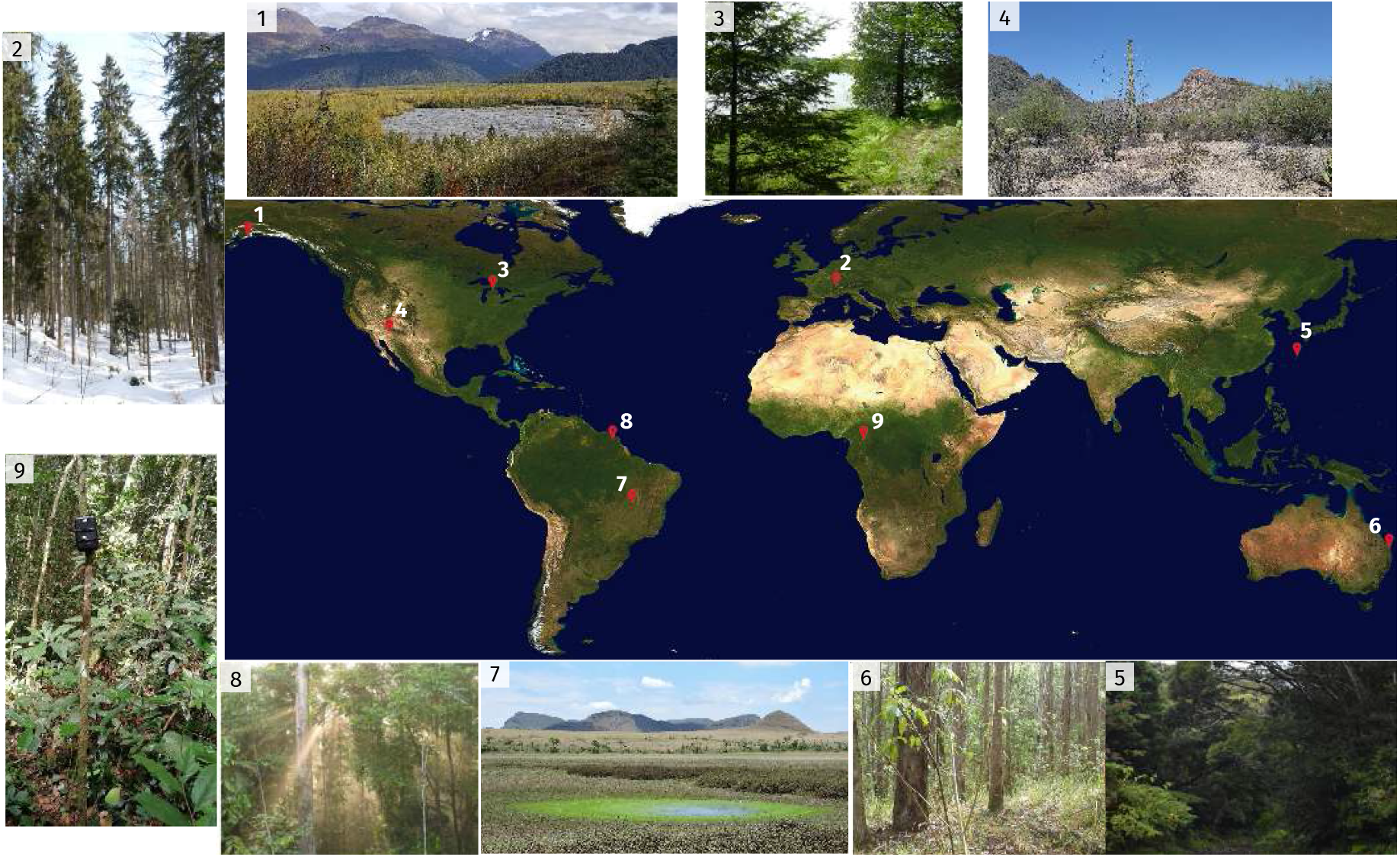
Pictures and locations of the nine habitats registered in the recordings from the Hearbiodiv database. Habitats’ numbers are sorted by distance to the Equator. 1: Kenai Fjords, Alaska, USA; 2: Haut Jura, Risoux, FRA; 3: Twin Lakes, Michigan, USA; 4: Ironwood, Arizona, USA; 5: Yanbaru, Okinawa, JPN; 6: Woondum, Queensland, AUS; 7: Chapada dos Veadeiros, Cerrado, BRA; 8: CNRS Research Site, Nouragues, GUF; 9: La Belgique Research Site, CMR. Details for each recording site are provided in table I

#### 2. Main factors

From all the available recordings a smaller selection was made, named the **Hearbiodiv database**, using a criteria that minimised the number of recordings to be analysed but preserved the heterogeneity of the soundscapes as much as possible. This heterogeneity is related, mainly but not exclusively, to patterns of biological activity and differences in the environment such as topography, vegetation, climate, etc. (Grinfeder *et al*., 2022b). Therefore, three main factors were considered to represent and summarise this heterogeneity in a way that was relevant for this study: habitat, moment of the day and precipitation period. Habitat is the most important factor as this study aims at characterising and distinguishing them. Moment of the day and precipitation period are the factors used to preserve the heterogeneity of the soundscapes in the Hearbiodiv database. In summary, four moments of the day and two periods of the year contrasting in precipitation level were selected for each habitat, considering also the limitation of the available recordings. The four moments of the day (midnight, sunrise, midday and sunset) were defined as the two hours closer to sunrise and sunset time on each day, and the two hours closer to the middle points between them for midnight and midday. The two periods of the year, named as “wet” and “dry”, were defined as the two months closer to the highest and lowest probability of precipitation respectively for each habitat.

##### Precipitation period

climatic information was extracted from the website `weatherspark.com` (owned by Cedar Lake Ventures, Inc. in USA) which gathers climatic information from different sources from several years. This information was used to define the precipitation period as wet or dry for each habitat. For each location, the closest city with climatic information available was selected (names specified in second row of table II). In particular, the information related to the probability of precipitation per month was used. This probability was computed by the website as the percentage of days in which various types of precipitation were observed (rain alone, snow alone, and mixed). Using this probability, the website defined the wet and dry periods for each location. Combining this information with the recordings available for each location allowed the selection of two months from each precipitation period. Whenever available, these months were consecutive for each period and as close to the wettest/driest months as possible, final selected months for each habitat and period are shown in table II.

**TABLE II.**
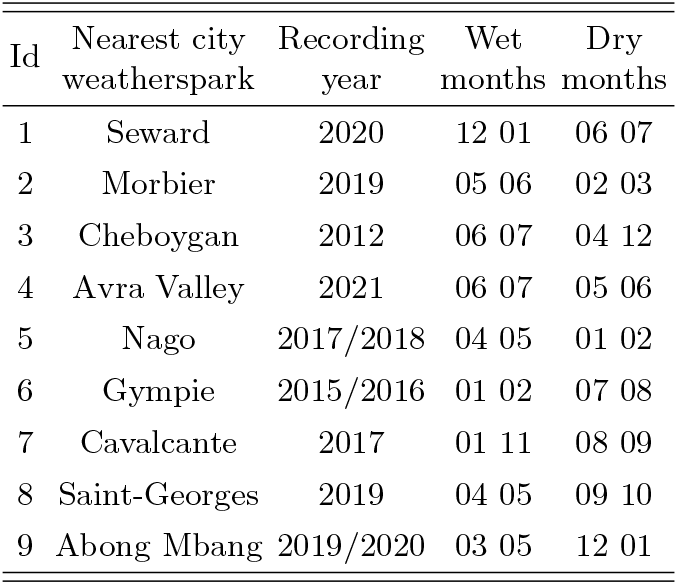
Specifications for selection of wet and dry precipitation periods.

##### Moment of the day

to define the four different moments of the day, sunrise and sunset times were computed for each selected day. These times were computed using the Matlab implementation by Droste (2022) of the sunrise/sunset and solar position calculations developed in excel/open office by the USA National Oceanic and Atmospheric Administration, based on equations from Astronomical Algorithms by Meeus (1991). With this implementation, the sunrise and sunset time obtained were corrected by atmospheric refraction effects so that these times are closer to the real sunlight (when sky colour changes) than just the position of the sun (when sun crosses the horizon line); the daylight saving time change is also taken into account when needed as UTC corrections for the corresponding months (UTC is an input for the Matlab function). Because animal behaviour is related to sunlight, astronomical, nautical, and civil twilight times were computed to ensure that the selected 2-hour window for sunrise and sunset were within these limits so the dawn and dusk choruses were not missed.

Once sunrise and sunset were computed, midday was defined as the two hours closer to the middle point between sunrise and sunset, and midnight as the two hours closer to the middle point between sunset and sunrise. Whenever midnight fell in between days (i.e., from 23:00 to 01:00) the definition was forced into one day only (i.e., from 00:00 to 02:00) so the four moments were properly defined for each day selected. Finally, due to restrictions in the recorded hours, all moments were defined by precise ‘o’clock’ times. This means that whenever an hour computed was below or above X:30, it was approximated to X or X+1, respectively.

#### 3. Selection procedure for acoustic recordings and creation of Hearbiodiv database

Figure 2 summarises this selection procedure with a schema. In order to make a systematic selection of sound samples, ten equi-spaced days were selected for each month from each precipitation period. Therefore the Hearbiodiv database is made of 40 days total (10 days x 2 month x 2 precipitation periods) for each habitat. Then, moments of the day were computed for each day. From each hour of each moment of each day, 4 equispaced seconds were extracted. These are the acoustic samples that make the Hearbiodiv database. Whenever this systematic selection was not possible for a given case due to a lack of recordings, the closest possible candidate was selected. This selection procedure was conducted blindly without any previous knowledge of the content of each recording to avoid any particular bias into the final selection. The only recordings that were removed from the study were those that included human voices or aircraft sounds (aircraft sounds were removed only for habitats 2 and 6, the only ones with plane information available).

**FIG. 2.**
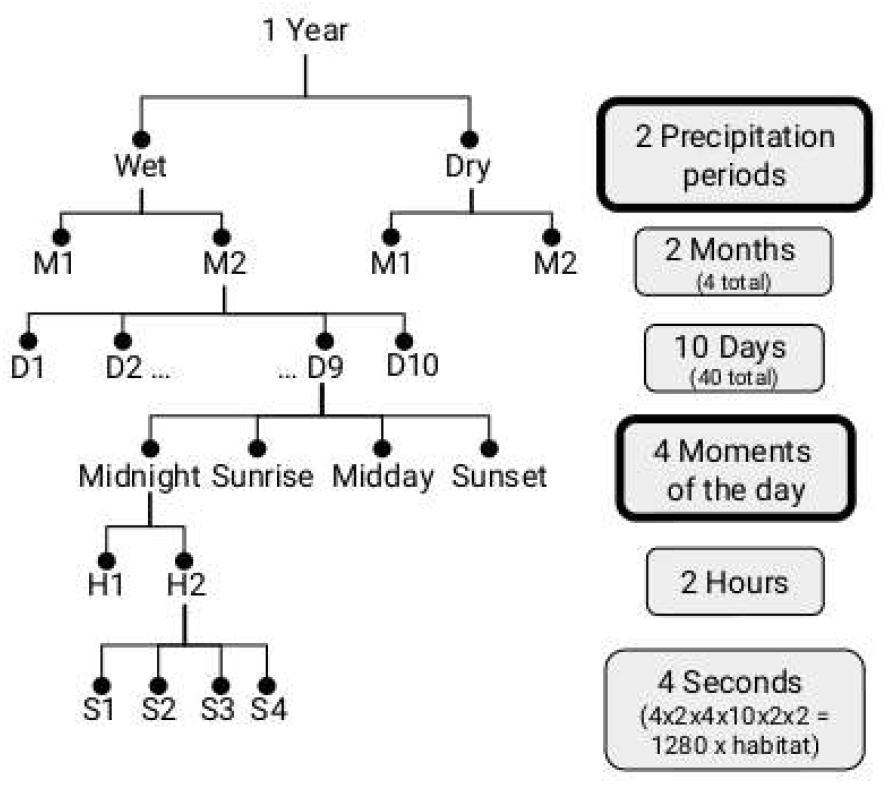
Schema that summarises the procedure for blindly selecting 1-sec long sound samples from available recordings.

The acoustic samples were limited to a duration of one second, so that they could be used in future behavioural protocols using the Hearbiodiv database (e.g., Apoux *et al*., 2023; Miller-Viacava *et al*., 2024; Thoret *et al*., 2020). A duration of one second limits the influence of cognitive factors (e.g., attentional and memory factors) in auditory detection, discrimination or recognition tasks involving modulation cues. A complementary analysis was conducted with 1-minute long samples to assess the effect of duration and is discussed in appendix A. The resulting final count of acoustic samples within the Hearbiodiv database is 1280 per habitat (2 precipitation periods x 2 months/period x 10 days/month x 4 moments of the day x 2 hours/moment x 4 seconds(samples)/hour = 1280 samples). This corresponds to 11520 1-sec long samples overall (1280 samples x 9 habitats = 11520 samples).

Finally, as the original acoustic recordings were made by different teams with different protocols, it was necessary to standardise all acoustic samples by using the lowest standard from the nine recording protocols. All 1-sec selected samples were (i) transformed into mono signals by choosing the right channel; (ii) downsampled so their sampling rate was fs=22050 Hz; (iii) normalised in long-term root mean square (RMS) power; and (iv) cosine ramps of 50 ms were applied to the beginning and end of each sample’s waveform.

The Hearbiodiv database used in the present study is available, depending on the project, upon request to the management.

### B. Computing the modulation power spectra (MPS)

#### 1. Modulation power spectra

Modulation power spectra (MPS) were computed for each 1-sec long sample to assess the spectro-temporal modulation information conveyed by the natural soundscapes used in this study. This MPS representation shows the joint spectral and temporal modulation power of sounds, that is the amount of energy (colour coded) as a function of temporal modulation frequency (x-axis) and spectral modulation frequency (y-axis) (Singh and Theunissen, 2003). It is obtained via a 2D short-time Fourier transform of the spectrogram of each acoustic sample. The frequencies obtained from the Fourier transform of the temporal axis of the spectrogram correspond to the x-axis of the MPS and show the temporal modulations within the sound (in Hz). The frequencies from the Fourier transform of the spectral axis of the spectrogram correspond to the y-axis of the MPS and show the spectral modulations within the sound (in cycles/kHz) (Elliott and Theunissen, 2009).

These MPS computations were done in Python using the library `BioSound` developed by Frederic Theunissen (Theunissen, 2018) which performs a gaussian windowed two-dimensional short-time fast Fourier transform (2D-ST-FFT) on the logarithmic amplitude of the spectrograms of the samples, which were computed via a gaussian windowed ST-FFT on the sound signals.

The final MPS representations used in this study are the logarithmic transformation of the amplitude output of the 2D-ST-FFT squared. By convention, only one of the MPS axes must have positive and negative frequency modulations to distinguish upward from downward frequency sweeps, and the temporal modulations is the commonly used one (e.g., Elliott and Theunissen, 2009). This logarithmic transformation is not included when computing modulation statistics.

### 2. Time-frequency scale and input parameters

The functions used within the `BioSound` library had several input parameters that were selected such that the final spectro-temporal resolution of the MPS was within the range of human perception: 0 *< ω*_*f*_ *<* 16 (cycles/kHz), −150 *< ω*_*t*_ *<* 150 (Hz) (Moore, 2013).

#### Spectrogram computations

using the function `spectroCalc`, the input parameters selected were `spec_sample_rate`=300, `freq spacing`=28.65, `min freq`=0, `max_freq`=None. Here `spec_sample_rate` is the desired sampling rate for the output spectrogram (in Hz), `freq_spacing` is the desired time-frequency scale for the spectrogram (in Hz), `min_freq` and `max_freq` define the interval of the frequencies to analyse. By setting `max_freq` to None the Nyquist frequency was used. These parameters determine within the `BioSound` code the width and overlap of the gaussian window as `window_length`=nstd/(2*π* `freq_spacing`)=0.033 sec (where nstd^2^ is the variance of the gaussian with nstd=6 default value used) and `overlap`=1-(24*π* `freq_spacing`/nstd*`spec_sample_rate`)=0.79 so 79% of the window with the parameters used. The 1-sec long samples were zero padded at their beginning and end by half of the window length each before any computations were done. All this results in spectrograms with 368 frequency bands between 0 and 11 kHz (with 30 Hz of spacing), and with a real sampling rate of 297 Hz, meaning 297 time data points for the 1-sec long samples (with 0.0034 sec of spacing).

#### MPS computations

calculated with the function `mpsCalc` which uses an overlap and add method with a gaussian window. The input parameters for this function were `window`=0.5, `Norm`=False. Here, `window` is the length of the gaussian window (in secs) and `Norm` controls for Z-score normalisation which is not used if set to False. These windows also have a variance of nstd^2^ = 12. After averaging the 2D-ST-FFT results for these windows, the MPS representations have a range of spectral modulations between 0 and 16.58 cycles/kHz divided into 184 segments spaced by 0.091 cycles/kHz, and a range of temporal modulations between -147.99 and 147.99 Hz divided into 150 segments spaced by 2 Hz.

### 3. Modulation statistics of MPS

Following the work of Singh and Theunissen (2003), six quantifiers were computed from the MPS to better describe and represent some of its structure. These quantifiers are referred to as the modulation statistics in this article.

#### Separability (*α*_sep_)

the MPS is defined as separable if the joint spectro-temporal modulations can be predicted from the mean spectral or mean temporal modulations estimated separately. Therefore, it assesses the degree of independence between spectral and temporal modulations in the stimulus. A higher degree of dependency (i.e., low separability) would predominantly come from the presence of several sound sources and/or frequency sweeps in the sound, which would result in more complex structure of the MPS (a completely random noise sound, which modulations are totally independent, shows the simplest possible structure of its MPS, and thus, the most separable one). Considering the MPS as a matrix, *α*_sep_ was estimated using the singular values (*λ*_*i*_) obtained from its singular value decomposition (SVD), because higher separability is reached for higher predominance of the first singular value (*λ*_1_) compared with the remaining ones (Depireux *et al*., 2001). Therefore

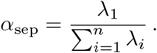

A maximum *α*_sep_ value of 1 indicates that the MPS is fully separable.

#### Asymmetry (*α*_asym_)

this statistic highlights the difference in modulation power between the left (*P*_up_) and right (*P*_down_) quadrants of the MPS. *α*_asym_ is estimated by computing the ratio

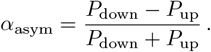

The amount of modulation power in the left quadrant relates to the amount of frequency up-sweeps in the sound, conversely, the right quadrant relates to the presence of down-sweeps. The higher *α*_asym_ is, the more downsweeps than up-sweeps the sound has, negative values being associated with a higher presence of up-sweeps than down-sweeps. A value of 0 means that the MPS is perfectly symmetric, meaning that the sound has an equal amount of frequency down-sweeps and up-sweeps.

#### Low-pass coefficient (*α*_low_) and Starriness (*α*_star_)

both of these statistics are related to the distribution of modulation power within the MPS and how uniform it is. One of the most remarkable findings of Singh and Theunissen (2003) is that human and non-human animal vocalisations exhibit the highest spectral modulation power only at slow temporal modulations. Low-pass coefficient highlights how much of the total modulation power (*P*_total_) is restricted in a low-pass region (*P*_low_) defined by slow temporal (−10 to 10 Hz) and low spectral modulations (0 to 0.195 cycles/kHz). So *α*_low_ is defined as the ratio

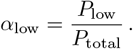

Starriness quantifies how much of the total modulation power that is outside the low-pass region is concentrated across both temporal and spectral axes of the MPS. This concentration across the axes highlights how much of the total modulation power (*P*_total_) is restricted to either slow temporal modulations (−10 to 10 Hz) but with a range of higher spectral modulations (0.195 to 16.576 cycles/kHz) 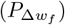 (for example, such a combination would appear for tonal/harmonic sounds), and/or low spectral modulations (0 to 0.195 cycles/kHz) but with a range of faster temporal modulations (10 to 147.987 cycles/kHz) 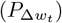 (for example, such a combination would appear for transient sounds like barks or insect stridulations). So *α*_star_ is defined as the ratio

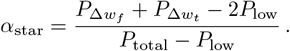

#### Modulation depth (*α*_mod_)

Modulation depth shows the overall strength of the modulations. It is estimated as a proportion of the modulation power at 0 temporal and spectral rates (*P*_dc_) in comparison to all other temporal and spectral modulation rates. Therefore, *α*_mod_ is the ratio

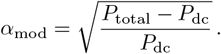

This modulation statistic is computed without the logarithmic transformation of the amplitude of the spectrograms.

#### Alpha (*α*_temp_)

this statistic conveys information about the shape of the temporal modulation spectrum only. It was obtained from fitting a power law relationship 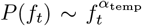 to the temporal modulation power spectrum. This statistic was computed with a different procedure than the one used by Singh and Theunissen (2003) as it capitalises on the SVD procedure for computing Separability and it was not known in advance whether natural soundscapes would show high Separability. Instead, it was done without the MPS representation but following the procedure of Varnet *et al*. (2017) for computing the so-called AMa spectrum. Varnet *et al*. (2017) compute the temporal component of the amplitude-modulation spectrum with an algorithm that (i) decomposes the audio signal (with a Gammatone filterbank) into its different frequency components, then (ii) extracts the temporal distribution of modulation power as a function of each frequency channel to finally (iii) compute the Fourier transform of these resulting envelopes to obtain the temporal-modulation frequency. The final AMa spectrum is the average of the output of the Fourier transform across all audio frequency channels given by the Gammatone filterbank. The power law relationship described above was fitted to this spectrum within a range of temporal-modulation frequencies between 3 and 100 Hz, which is the same range used by Singh and Theunissen (2003) in their fitting procedure.

### C. Data analysis

Modulation statistics defined above were analysed mainly with a non-parametric aligned rank transform (ART) three-way ANOVA (using ARTool toolbox in R by Wobbrock *et al*., 2011) to assess the influence of habitat, moment of the day, and precipitation period. More details concerning this statistical method are provided in appendix B. A full pairwise comparison post-hoc test was conducted (using ART-C contrasts within the AR-Tool in R by Elkin *et al*., 2021) with a conservative Holm–Bonferroni adjustment of *p*-values for significance due to multiple comparisons. These results are detailed in section III A.

Additional analyses were later implemented to further expand on the results obtained from different perspectives as detailed below.

#### 1. Habitat classification

To assess the possibility of using modulation information (as summarised by modulation statistics) for distinguishing habitats, results from the pairwise comparison described above were analysed by habitat. This is discussed in section III B. In addition, a machine learning classifier was trained to distinguish the nine different habitats from the Hearbiodiv database using the modulation statistics. Moreover, the contribution of each modulation statistic to habitat classification was assessed by analysing the feature importance ranking. This machine learning analysis was done on python with `sklearn` and `xgboost` libraries. Results from this classifier are discussed in section III C.

##### Model selection

An extreme gradient boosting (XGBoost) algorithm was chosen as it is one of the most powerful models and has shown high performance in a wide range of tasks. Multiclass classification error rate (merror) was used to train the model. Model performance was evaluated using the number of correctly classified samples (i.e., the accuracy score).

The data, corresponding to all modulation statistics values, were pre-processed before the model fit procedure. Given the nature of the data, however, a search for outliers was not performed. By construction, the data does not include missing or incorrect values. Therefore, preprocessing simply consisted in z-score normalisation of the features, which correspond to the six different modulation statistics.

The model was used to classify the data according to their original habitat. As mentioned previously, the database included 1280 samples for each of the nine distinct habitats. All samples were used and randomly split into two groups, with 80% of the data (1024 samples) designated as the training set and the remaining 20% (256 samples) as the test set.

##### Feature importance

The set of six modulation statistics was evaluated for importance using three different approaches:

i. Mean decrease impurity: importance was first calculated using the built-in `feature_importance` attribute. This procedure measures, for each node in a tree, how the feature used for this split decreases the “impurity” (the default Gini impurity metric was used here). This is done for all the trees and the average decrease in impurity is calculated for each feature. The average over all trees in the forest is the measure of the feature importance.
ii. Permutation-based feature importance: importance was also calculated using the `permutation_importance` function from the `Scikit-learn` library. This permutation method randomly shuffles each feature and computes the difference between the baseline metric and the metric from permuting the feature column. The features which impact performance the most are defined as the most important.
iii. Feature selection: a last approach was implemented to corroborate the feature importance results obtained using the previous techniques. This approach involves training a model with a selected subset of features, and then evaluating this new model. A common procedure was used consisting of training a model while removing one feature at a time. However, not all combinations were evaluated. In this procedure, features were removed from least to most important (based on mean decrease impurity). Therefore, a first model was trained and evaluated with all 6 features, then the least important feature was left out, the second least important, etc., until a last model was trained and evaluated with the most important feature only. The importance of each feature can be deduced from the drop in accuracy observed when this particular feature is removed from training.

#### 2. Contributions of biophony and geophony

The final additional analyses concerns the exploration of potential contributions of biophony and geophony, independently, to the information captured by the six modulation statistics. For this, additional MPS analyses were conducted on recordings of sounds from different predominant sources (referred to as isolated sounds even though they are not all truly isolated (clean) recordings) and artificial mixtures of these sounds. Mixtures were created for biophony sounds only in order to investigate how a chorus of sounds made by living beings modified modulation statistics compared to isolated sounds. This is of particular interest as large animal choruses are expected to be more frequent in habitats with higher acoustic biodiversity. These additional analyses are discussed in section III E.

##### Isolated sounds

Ten different isolated sound sources were used divided in three big categories (birds, other animals and geophony). The birds category is composed of three groups with sounds from different locations, these are: (i) Birds HJ: 86 cleaned bird samples with no background sounds from Haut Jura, taken from 8 different species (provided by Jérôme Sueur, MNHN); (ii) Birds NG: 36 bird samples within their soundscape from Nouragues, taken from 5 different species (provided by Jérôme Sueur, MNHN); (iii) Birds YB: 22 bird samples within their soundscape from Yanbaru, taken from 7 different species (provided by Nicholas Friedman). The other animals category is composed of three groups with sounds from different sources but from the same location, these are: (i) Frogs, (ii) Insects and (iii) Mammals. They have 12, 14 and 35 biophony samples respectively, within their soundscape from Nouragues, taken from 2 different species for each source (provided by Jérôme Sueur, MNHN). Finally, the geophony category is composed of four groups with sounds from different sources and different locations (none of them part of the Hearbiodiv database), these are: (i) Rain: 7 samples taken from 1 recording; (ii) River: 17 samples taken from 3 different river recordings; (iii) Thunder: 13 samples taken from 1 recording; (iv) Wind: 50 samples taken from 10 different wind recordings (all geophony recordings provided by Richard McWalter except one river sound provided by Bernie Krause, Wild Sanctuary).

All isolated sound samples were cut to 1-sec long and resampled (fs=22050) to have the same parameters as the samples from the Hearbiodiv database.

##### Artificial mixtures

Four categories of mixtures were done, each with animals from different species but from the same location. 3 categories correspond to mixtures of birds (Birds HJ, Birds NG, Birds YB) and 1 (Mix NG) to mixtures of frogs, insects and mammals. All bird mixtures were made as combinations of 6 species, one sample from each. Mix NG were combinations of 2 frogs, 2 insects and 2 mammals species, one sample from each. Birds HJ: 6 mixtures; Birds NG: 2 mixtures; Birds YB: 2 mixtures; Mix NG: 3 mixtures.

## III. RESULTS

### A. MPS and modulation statistics across habitats, moments of the day and precipitation periods

Figure 3 shows MPS averaged from 160 acoustic samples for each habitat, moment of the day, precipitation period combination. These plots do not show the full range of spectro-temporal modulation rates explored here but zoom into the region with highest modulation power and, therefore, the most informative part of the MPS. Overall these plots show that regardless of habitat, moment of the day and precipitation period, most of the modulation power is concentrated in the central region of the full range of spectro-temporal modulation rates explored in this study. This means that natural soundscapes are characterized by low spectral (<0.5-1 cycles/kHz) and slow temporal (<10-20 Hz) modulations.

**FIG. 3.**
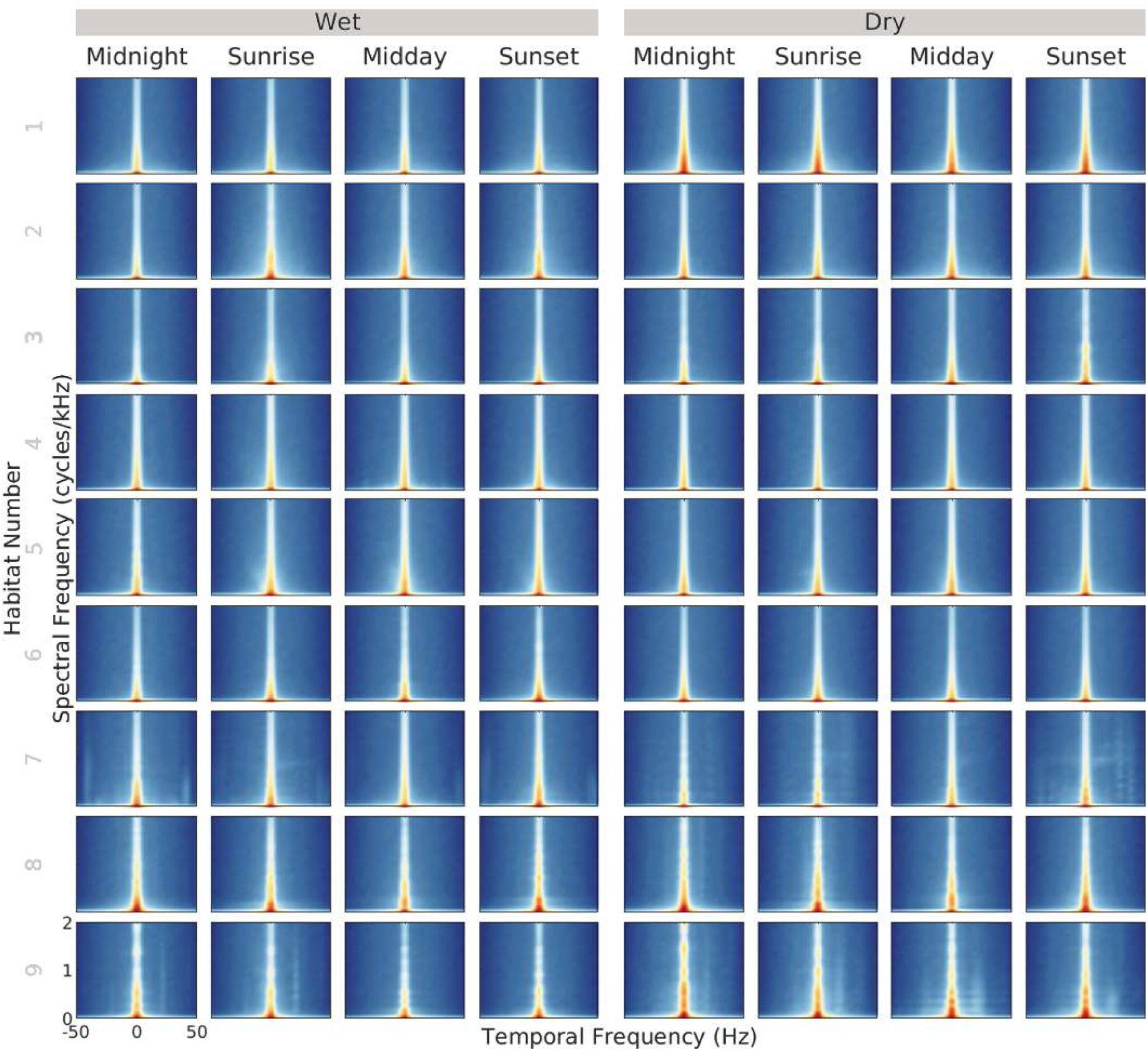
MPS for each habitat (one row each) sorted from 1 to 9 by their distance to the Equator (|latitude| of each location in degrees) as shown in table I. Separated by precipitation period (left panels: wet, right panels: dry) and moment of the day (one column each). Results averaged from 160 acoustic samples for each habitat, moment of the day, precipitation period combination (1280 total samples per habitat, 11520 samples overall). These plots do not show the full range of spectro-temporal modulation rates explored but a zoom into the regions with highest modulation power and, therefore the most informative.

However, within this range of predominant modulation power concentrated in the lower modulation rates, some differences appear between habitats, moment of the day and precipitation period combinations. For example: (i) habitat 3 and 8 show different structures in the distribution of modulation power (8 showing a more sinusoidal shape than 3); (ii) habitat 9 (which structure is also rather sinusoidal compared to other habitats) has its bumps distributed in different ways across different moments of the day; (iii) modulation power of habitat 1 encompasses a larger range of spectral modulations for the dry period compared to the wet period. The observed differences in modulation power are not consistent across habitats, moment of the day and precipitation period so there does not seem to be a specific combination of spectro-temporal modulation rates that would emerge as a marker of the effects or contributions of different habitats, moment of the day or precipitation periods. Still, the differences in modulation power seem to indicate that the modulation information available may be different across habitats, moment of the day and precipitation period combinations, and that this information may potentially be used to characterise and distinguish them.

Moving beyond the MPS representation and into modulation statistics, the distributions of all six modulation statistics (from all sound samples in the Hearbiodiv database) are presented in figure 4 for all nine habitats. These results are separated by precipitation period and moment of the day, which means a total of 72 plots for each modulation statistic (72 combinations of the three factors studied: 9 habitats x 4 moments of the day x 2 precipitation periods). These plots show that modulation statistics do not always follow a normal distribution, which was further assessed with QQ-Plots and Shapiro-Wilk tests applied directly to the distributions (this is discussed in appendix B). Moreover, the shape of the distributions differs across habitats, moments of the day and precipitation periods.

**FIG. 4.**
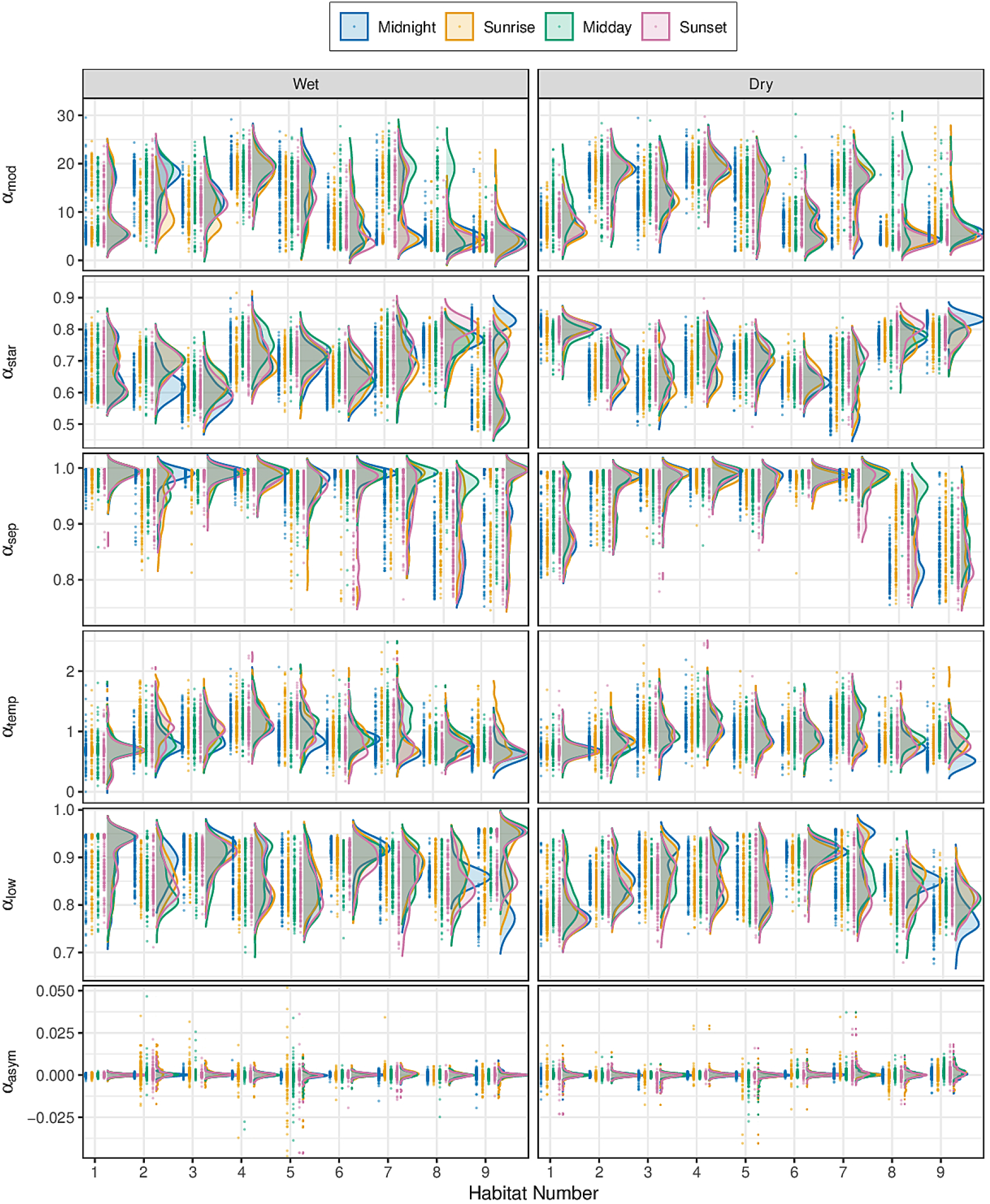
From top to bottom: data (points) and density distribution estimations (lines and surfaces) of all six modulation statistics extracted from the MPS, plotted for each habitat sorted from 1 to 9 by their distance to the Equator (|latitude| of each location in degrees) as shown in table I. The distributions are separated by precipitation period (left panels: wet, right panels: dry) and moment of the day (one colour/line type for each moment). Results from 160 acoustic samples for each habitat, moment of the day, precipitation period combination (1280 total samples per habitat, 11520 samples overall).

Figure 5 shows the means of all six modulation statistics extracted from the MPS, for all nine habitats sorted by their distance to the Equator (|latitude| of each location in degrees). These results are separated by precipitation period and moment of the day. In general, almost all modulation statistics values change across habitats for all moments of the day and precipitation periods. They show complex variation patterns with different trends across habitats, moment of the day and precipitation periods. More details on these trends and their possible causes are discussed below at the end of the Results section in III F.

**FIG. 5.**
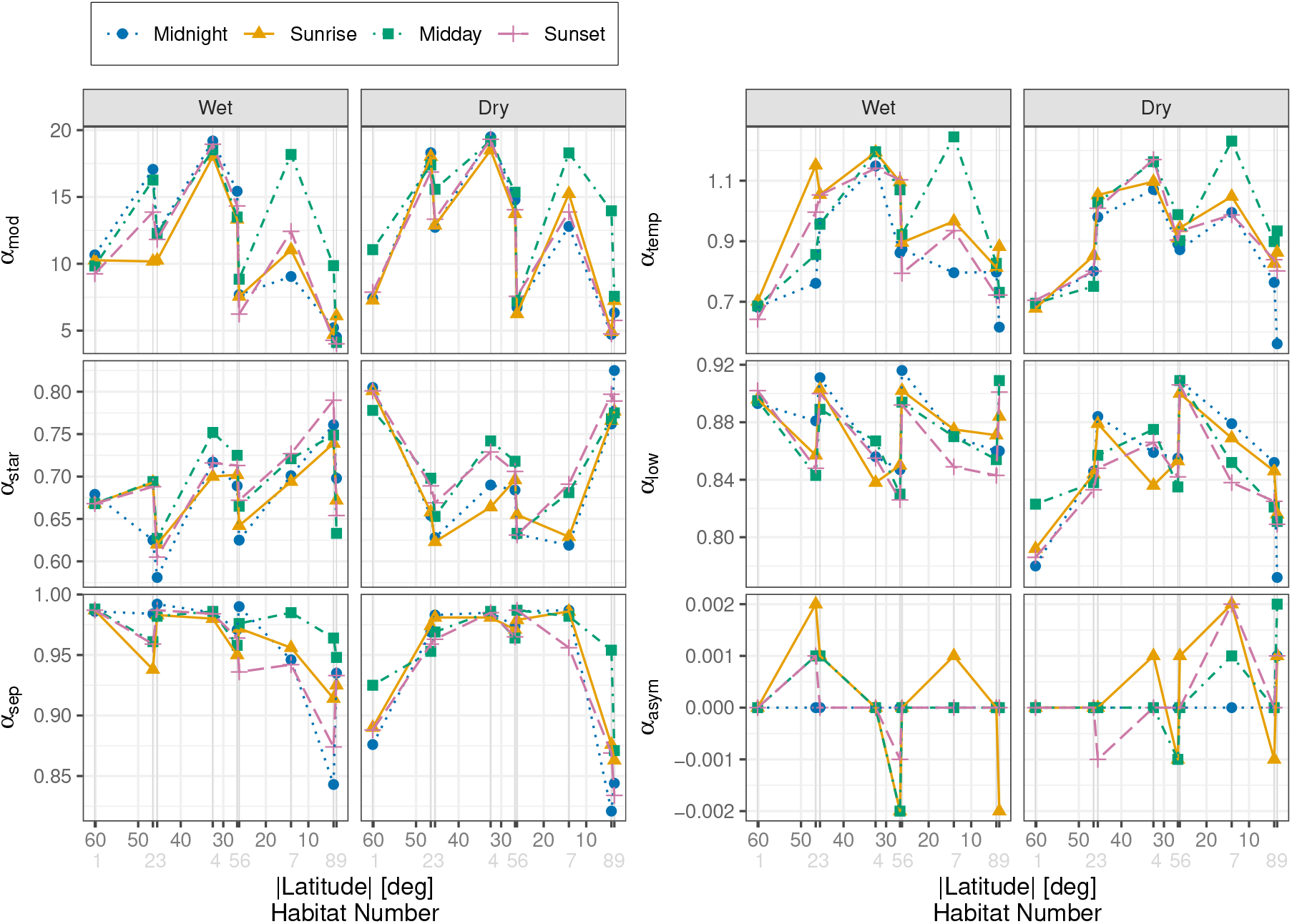
Mean of all six modulation statistics extracted from the MPS for all habitats, plotted as a function of each habitat’s distance to the Equator (|latitude| of each location in degrees) as shown in table I. Results for each statistic are separated by precipitation period (wet or dry panels) and moment of the day (one colour/line type/point type for each moment).

As discussed in section II C, differences across modulation statistic means observed in figure 5 were analysed by conducting a non-parametric aligned rank transform three-way ANOVA to assess the influence of habitat, moment of the day, and precipitation period on each modulation statistic. Significant three-way interactions were found for all modulation statistics as shown on table III. To check if the differences observed in modulation statistics across habitats depend on both moment of the day and precipitation periods, two separated additional test of two-way interactions were conducted, one for Habitat:Moment and another for Habitat:Period interactions. These analyses showed statistically significant two-way interactions between habitat and moment for all periods and also between habitat and period for all moments (all *p*-values ≤ 0.008, the significance level after Bonferroni correction: 0.05 divided by the number of two-way interaction computed (6)). This means that differences in all modulation statistics across habitat significantly depend on both moment of the day and precipitation period.

**TABLE III.**
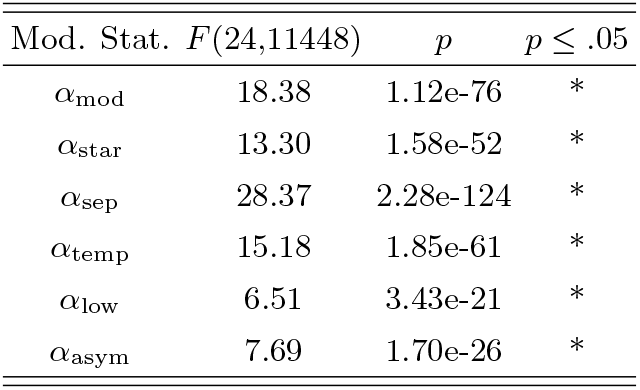
Significant three-way interactions (Habitat:Moment:Period) resulting from all six ART ANOVAs, one for each modulation statistic.

| Mod. Stat. | $F(24,11448)$ | $p$ | $p \leq .05$ |
| --- | --- | --- | --- |
| $\alpha_{\text{mod}}$ | 18.38 | 1.12e-76 | * |
| $\alpha_{\text{star}}$ | 13.30 | 1.58e-52 | * |
| $\alpha_{\text{sep}}$ | 28.37 | 2.28e-124 | * |
| $\alpha_{\text{temp}}$ | 15.18 | 1.85e-61 | * |
| $\alpha_{\text{low}}$ | 6.51 | 3.43e-21 | * |
| $\alpha_{\text{asym}}$ | 7.69 | 1.70e-26 | * |

Since all interactions are significant, to further study the influence of all potential combinations of the three factors studied without any a priori assumptions or hypotheses, a full pairwise comparison post-hoc test was conducted (details in section II C). All combinations of the three factors make a total of 2556 pairs analysed for each modulation statistic (a pair is formed by 2 random selections of any of the 72 possible combinations of the three factors without repetition 72×71 /2 = 2556). The significance of these comparisons are summarised in plots (one for each modulation statistic) presented in the first and third columns of figure 6. These plots show colour coded *p*-values for all possible pairings sorted by habitat. A first global result can be extracted from these comparisons by computing the percentage of total amount of pairs that show significant differences for each modulation statistic, which is shown between parentheses in the title of each modulation statistic on figure 6 and on the first row of table IV. Overall, the total amount of significant pairings varies across modulation statistics with Modulation depth being the one with the most significant differences and Asymmetry the lowest. This means that Modulation depth (79%) is the modulation statistic that captures most variability across habitat, moment of the day and precipitation period, followed by Starriness (78%), Separability (71%), Alpha (70%), Low-pass coefficient (67%) and Asymmetry (20%).

**TABLE IV.** Percentage (%) of total amount of pairs with significant differences from pairwise comparison post-hoc analysis for each modulation statistic. Summary for habitats. Highlighted all values ≥ 79%.

| Habitat | $\alpha_{\text{mod}}$ | $\alpha_{\text{star}}$ | $\alpha_{\text{sep}}$ | $\alpha_{\text{temp}}$ | $\alpha_{\text{low}}$ | $\alpha_{\text{asym}}$ |
| --- | --- | --- | --- | --- | --- | --- |
| All | 79 | 78 | 71 | 70 | 67 | 20 |
| 1 | **80** | **79** | **80** | **81** | **80** | 10 |
| 2 | **79** | 72 | 66 | 67 | 59 | 13 |
| 3 | 76 | **83** | 67 | 67 | 68 | 20 |
| 4 | **81** | 78 | 65 | 77 | 59 | 10 |
| 5 | 75 | 72 | 66 | 65 | 59 | 27 |
| 6 | 77 | 77 | 66 | 61 | 74 | 12 |
| 7 | 77 | 74 | 68 | 70 | 61 | 36 |
| 8 | **82** | **86** | **85** | 66 | 61 | 14 |
| 9 | **81** | **80** | 76 | 75 | **79** | 38 |

**FIG. 6.**
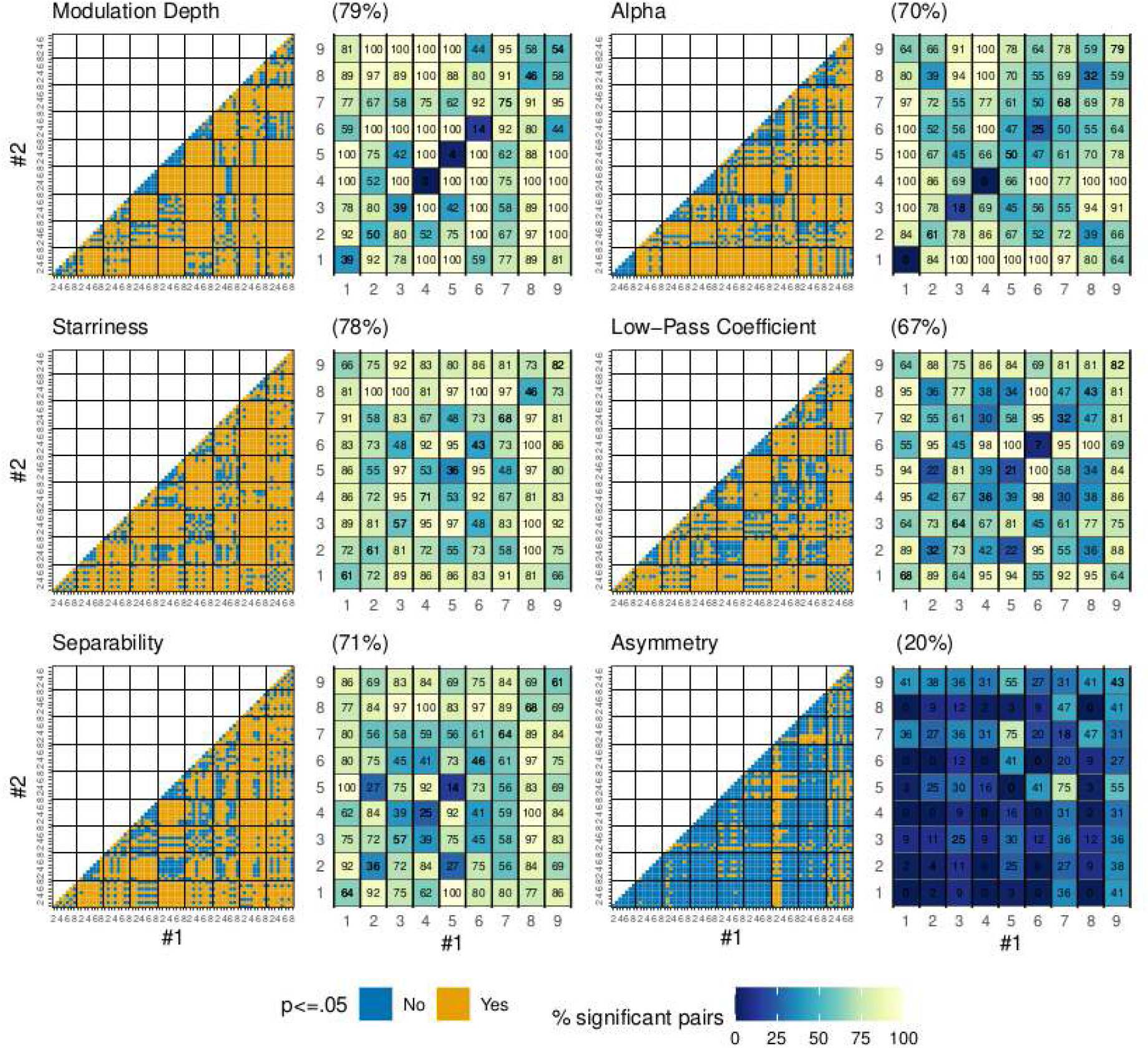
Pairwise comparison for ART ANOVA for all six modulation statistics. First and third columns: Significance of pairwise comparisons with Holm-Bonferroni adjustments of *p*-values. The x-axis shows one of the 72 possible combinations and the y-axis the second one needed to form a pair (a given combination is not compared with itself). Data in both axes is sorted by habitat from 1 to 9. Numbers 1 to 8 in tick marks of both axes represent the 8 possible combinations of moment of the day and precipitation period (odd numbers dry season, even numbers wet season, 1 to 4 twilight moments of the day, 5 to 8 midday and midnight: 1-sunrise,dry; 2-sunrise,wet; 3-sunset,dry; 4,sunset-wet; 5-midday,dry; 6-midday,wet; 7-midnight,dry; 8-midnight,wet). Second and fourth columns: Pairwise comparison summary for habitats. Percentage of pairs with significant differences for each habitat combination (28 pairs for within habitat comparison in the diagonals, 64 pairs across habitat). The x-axis shows one of the 9 possible habitat and the y-axis the second one needed to form a pair. Titles show in parenthesis the percentage of total pairs with significant differences found for each modulation statistic.

### B. General trends for habitat classification

Focusing the analysis on the capacity of the modulation statistics to characterise and distinguish habitats, a second result can be obtained from the full pairwise comparison by computing the percentage of total amount of pairs that show significant differences for each habitat, considering all possible combinations of moments of the day and precipitation periods. These results are shown in the rows numbered from 1 to 9 (one row for each habitat) of table IV. Differences across habitats are not the same for each modulation statistic. Additionally, the habitats that appear to be the most different across modulation statistics are habitats 1, 8 and 9, which are the farthest and closest to the Equator.

Detailed comparisons across habitats can be seen on the second and fourth columns of figure 6 where percentage of pairs with significant differences are shown for each possible habitat combination, for all moments of the day and precipitation periods combinations. These plots further highlight that the differences and similarities across habitats are not the same for all modulation statistics. Additionally, the diagonals in these plots (within habitat comparison) show the effects of moment of the day and precipitation period on modulation statistics which, overall, remain lower than the effects of habitat.

Overall, these results suggest that the present set of modulation statistics captures relevant information corresponding to different characteristics of the nine habitats, which may allow to characterise and distinguish them.

### C. Machine learning habitat classification

To further unveil whether spectro-temporal modulation information summarised by all six modulation statistics can distinguish across habitats, a machine learning classifier was trained to distinguish the nine different habitats from the Hearbiodiv database using the modulation statistics. All details for this procedure are given in section II C.

Accuracy obtained without any optimisation was 65% (chance level = 11% when randomly choosing one out of nine possible habitats). Grid search was conducted to fine-tune some of the hyperparameters of the model (`max depth`, `eta` and `gamma`). However, the performance of the model did not improve markedly compared to the 65% accuracy score obtained using the default hyperparameter values. As a consequence, the latter values were used for easier reproducibility. A detailed performance of the classification model can be seen in the confusion matrix shown on figure 7 where predicted and real habitat labels are compared for the 256 sounds for each habitat. Information on the diagonal shows total of correctly classified habitats, which ranked from highest to lowest is (with percentages of total correctly classified sounds): 1 (81%), 8 (73%), 9 (73%), 6 (72%), 4 (66%), 2 (58%), 5 (58%), 3 (55%), and 7 (46%). Consistent with the statistical analysis shown on the previous section III B, habitats 1, 8 and 9, the farthest and closest to the Equator, seem to be the most differentiable from the nine habitats within the Hearbiodiv database, when using information from modulation statistics. Interestingly, habitat 7 is mostly confused with 4 (both are open habitats) and 9 with 8 (both are tropical habitats).

**FIG. 7.**
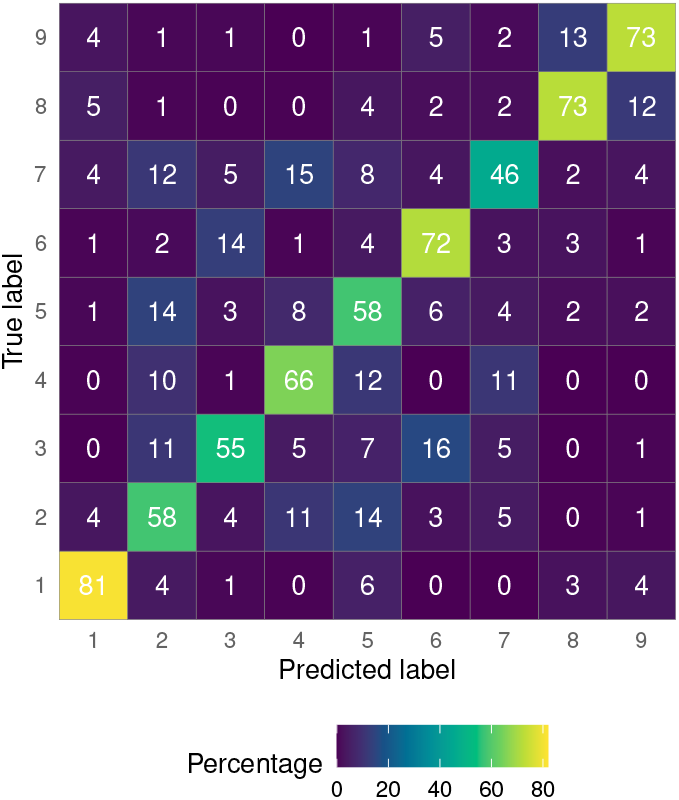
Confusion matrix for XGBoost computation of habitat classification. The matrix shows percentage of total sounds from each given habitat (true label) with respect to the habitat that they were classified as (predicted label). There were 256 sounds for each habitat.

With respect to feature importance, the first two evaluations detailed in section II C led to similar qualitative results. The first approach (mean decrease impurity) indicated that the ranking of importance of the features was as follows: Modulation Depth (0.26), Starriness (0.20), Separability (0.17), Alpha (0.15), Low-pass (0.14), and Asymmetry (0.07). The second approach (permutation) led to a slightly different order with an inversion of the two most important features: Starriness (0.26), Modulation Depth (0.24), Separability (0.23), Alpha (0.16), Low-pass (0.14), and Asymmetry (0.03). Finally, the performance of six models was assessed using the previously explained feature selection approach. Each model included the six, five, four, three, two, or single most important features, resulting in an overall accuracy of 65%, 63%, 60%, 54%, 42% and 30%, respectively. These results confirm the importance ranking obtained by measuring the mean decrease impurity, as removing the most important features has a strong effect on accuracy. These results are consistent with the statistical analysis shown on the previous section III B, as percentage of total significant pairings for modulation statistics follow the exact same rank (Modulation Depth, Starriness, Separability, Alpha, Low-pass, and Asymmetry).

Overall these results show that modulation statistics could be used to classify different habitats. Even without optimisation, the accuracy score of 65% remains well above chance level (11%). Moreover, and despite the minor discrepancy observed between the first two approaches, the results of the three feature importance assessments suggest that Modulation depth and Starriness are the two most important features and that Asymmetry has a very limited contribution to the decision-making process.

## D. Contributions of sound propagation: open versus closed habitats

All modulation statistics in figure 5 show complex variations that could be influenced by many factors that differ across habitats such as temperature variations, vegetation type and density, biodiversity, etc. From all these factors, vegetation plays an important role in altering sounds when they are transmitted through it. Habitats with high vegetation density are referred to as closed while the ones with low density as open. From the Hearbiodiv database, two are open (4 and 7) and seven are closed (1, 2, 3, 5, 6, 8, and 9).

Even if the number of open habitats was low and should be increased in further analyses, when comparing open and closed habitats in figure 5, habitats 4 (desert) and 7 (savannah, midday) may be distinguished from the remaining habitats due to their higher values of Modulation depth and to a lesser extent Alpha. These modulation statistics relate to total amount of modulation power, so it makes sense that these two statistics have the highest capacity of distinguishing open from closed habitats, as the effect of vegetation on sound propagation strongly impacts the spectro-temporal modulation content of the sounds as detailed below.

At least three main vegetation effects are known to impact sound propagation (see Forrest, 1994, for a review): (i) scattering and resulting filtering effects, (ii) reverberation effects, and (iii) atmospheric turbulence. These effects have different impacts on open versus closed habitats and alter transmission of modulation information differently. Excess in attenuation (re: spherical spreading) due to vegetation and resulting scattering should degrade modulation transmission of all temporal modulation rates and these effects should be stronger for closed than open habitats (Morton, 1975; Wiley and Richards, 1978). Reverberation is expected to affect specifically fast temporal modulation rates through smearing effects and the latter should be stronger in closed than open habitats (Wiley and Richards, 1978). Finally, atmospheric turbulence generates relatively slow random fluctuations patterns with more modulation power at relatively slow temporal-modulation rates, and these effects should be stronger for open than closed habitats (Waser and Brown, 1986).

Altogether, the specificities of sound propagation for natural settings may explain why soundscapes for the open and closed habitats of the Hearbiodiv database show distinct modulation characteristics, with open habitats showing more pronounced temporal-envelope fluctuations at relatively slow rates (i.e., higher values of Modulation depth → more modulation power → more spectro-temporal modulations in sounds overall, higher values of Alpha → steeper slope in temporal modulation spectrum → temporal-envelope fluctuations concentrate at slower rates).

## E. Contributions of biophony and geophony

In an attempt to disentangle the respective contributions of biophony and geophony to the spectro-temporal modulation information captured by the six modulation statistics, additional MPS analyses were conducted on isolated sounds and mixtures of these sounds. All details for this procedure are given in section II C. The six modulation statistics for isolated sounds and their mixtures are shown on figure 8.

**FIG. 8.**
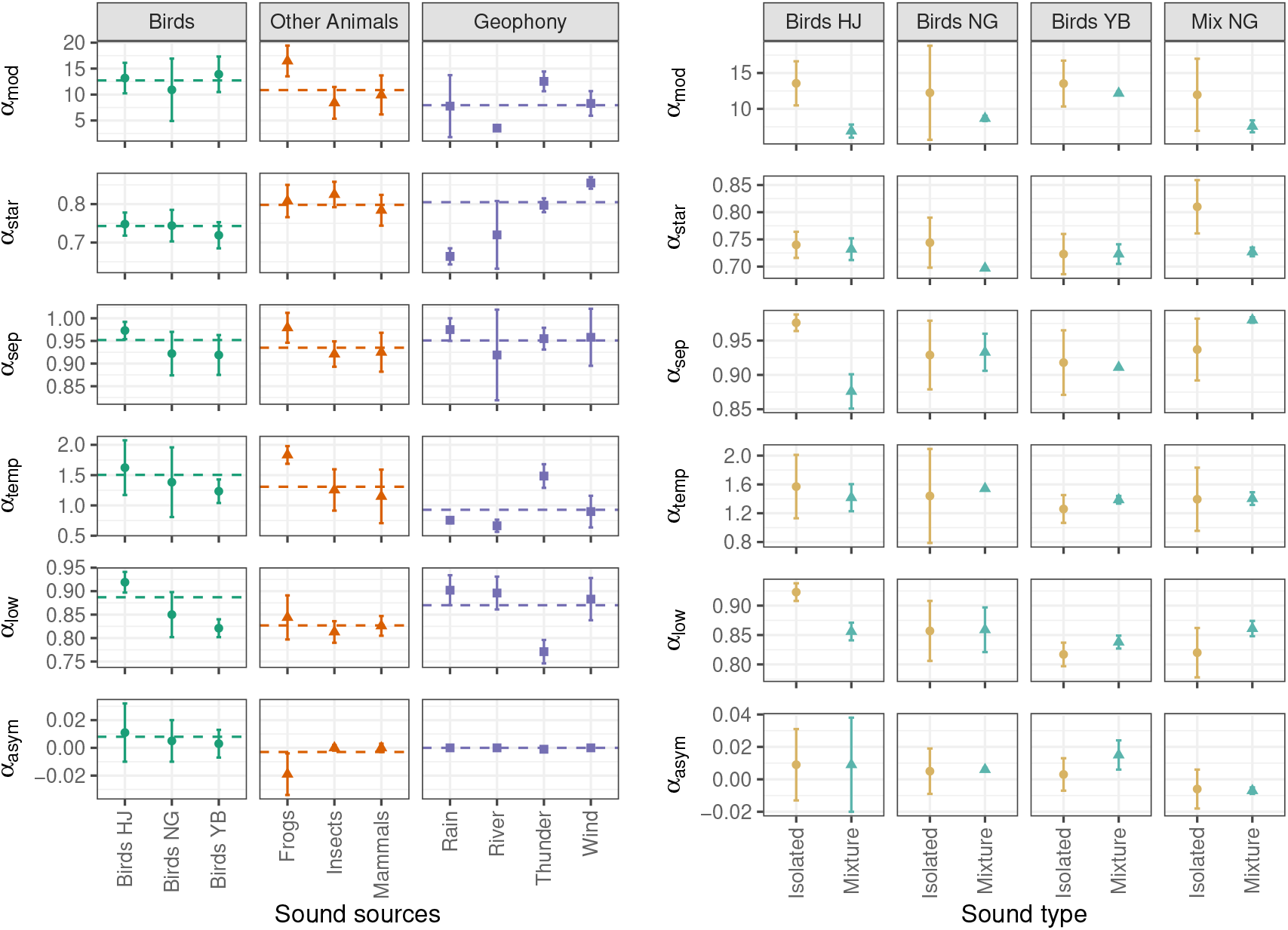
Mean and standard deviation (error bars) of each modulation statistic extracted from the MPS. Left: Ten different isolated sound sources divided in three categories (birds, other animals and geophony). Points show mean values for sounds in each group, dashed lines show means for each category. Right: Comparison between means of isolated sounds and mixtures created with these isolated sounds, divided in four groups depending on their sound sources. Details for isolated sounds used and artificial mixtures created are discussed in section II C.

Results from isolated sounds suggest that Modulation depth and Alpha have higher values for biophony than geophony, suggesting that overall, biotic sound sources (e.g., animal vocalisations) convey more salient modulation cues overall and their temporal-envelope fluctuations are more concentrated at slower rates. Asymmetry of geophony is generally lower than biophony (especially when compared to bird sounds), indicating that biophony conveys slightly more down-sweeps. These results are in concordance with previous findings by Singh and Theunissen (2003) who compared isolated recordings of Zebra finch songs with a collection of different geophony recordings (rustling brush, rain, fire, and stream sounds amongst others). However, comparison with their results is limited as the characteristics of the sounds are very different compared to the ones used here (e.g., Singh and Theunissen (2003) only studied a single bird species (Zebra finch) recorded under totally isolated conditions, and their geophony recordings were much longer). In any case, it is interesting to notice that results for the remaining modulation statistics (Starriness, Separability and Low-pass coefficient) are not consistent with the ones obtained by Singh and Theunissen (2003) who found that Starriness and Low-pass coefficient is lower for environmental sounds than Zebra finch songs, and Separability is slightly higher. The reasons behind these discrepancies remain unknown. An in-depth study based on a larger database is warranted to further explore these differences. Still, the current results suggest that rain and stream sounds might show lower Starriness than biophony. Moreover, variability of Starriness values is higher within geophony than biophony which suggests that spectro-temporal modulations of biophony are more constrained than geophony. This difference between biophony and geophony may presumably be due to strong constraints in vocal production of animals (i.e., they cannot produce rapid temporal modulation and high spectral modulations simultaneously).

Results from sound mixtures are easier to interpret for the “Birds HJ” group as these bird sounds have been highly filtered to remove as much background sound as possible. This means that the differences observed for this group’s mixtures are driven mainly by the effect of adding more animal sounds. These results indicate that an animal chorus has lower Modulation depth, Separability and Low-pass coefficient values compared to isolated animal sounds. These differences are lost in the “Mix NG” group for Separability and Low-pass coefficient, who show an opposite behaviour. In addition, this group shows lower values of Starriness for the mixtures compared to isolated sounds. Differences between both sound types in this group can be attributed to either the mixture of the main sound sources, the residual background sounds, or a combination of both. Disentangling the contribution of each would require an in depth study that is beyond the scope of this research.

## F. General trends for modulation information and possible causes

To summarise all main results discussed previously, the analyses conducted with the Hearbiodiv database show that:

- Spectro-temporal modulation power of natural soundscapes is mainly concentrated at slow temporal modulations (<10-20 Hz) and low spectral modulations (<0.5-1 cycles/kHz).
- Modulation statistics capture relevant information from habitats that have the potential to characterise and distinguish them.
- Modulation depth, Starriness, Separability and Alpha show the highest variations overall when analysing natural soundscapes. In particular, they proved to be the most relevant when classifying different habitats. In contrast, Asymmetry is the least informative for habitat classification as it shows the lowest variations overall when analysing natural soundscapes.
- Although still significant, effects of moment of the day and precipitation period on modulation statistics are lower compared to effects of habitat.
- Modulation statistics overall are particularly effective (i.e., they show the highest significant differences) at distinguishing habitats 1, 8 and 9 (the farthest and closest to the Equator) from the remaining ones.
- Due to sound propagation effects, Modulation depth and Alpha are potentially good candidates to distinguish open from closed habitats (higher values being found for open than closed habitats).
- Modulation depth, Alpha and to a lesser extent Asymmetry, are potentially good candidates to distinguish biophony from geophony within natural soundscapes (higher values being found for biophony than geophony).
- The range of Starriness values is higher for geophony than biophony, suggesting that spectrotemporal modulations are more constrained for biophony.
- Modulation depth, Separability and Low-pass coefficient are higher for isolated sound sources than animal choruses.

These results suggest some possible interpretations for the patterns observed in figure 5:

i. **Modulation depth** shows some of its highest values for habitats 4 and 7 (midday) and lowest for 8 and 9. For habitats 4 and 7, this may be due to their low density vegetation (open habitats) while habitats 8 and 9 combine effects of both having higher density vegetation (closed habitats in the tropical region) and higher biodiversity (presumably due to the biodiversity gradient as they are the closest to the Equator), and both effects make overall modulation depth lower. In between values of modulation depth (habitats 1, 2, 3, 5 and 6) are harder to interpret as contrasts of biophony/geophony and isolated/choruses sound sources show opposite effects (it seems that animal choruses may bring modulation depth values low enough to make them comparable to the ones of soundscapes dominated by geophony), and additionally, effects of slightly different vegetation density are harder to quantify. Presumably, habitats dominated by biophony but with more predominant isolated songs should have higher modulation depth than the ones with predominant animal choruses.
ii. Regardless of habitat, precipitation period, and moment of the day, **Low-pass coefficient** remains above 77% and **Starriness** above 67% for all the sounds studied. This indicates that overall at least 77% of modulation energy remains concentrated in the low-pass region, and from the remaining 23% outside, more than half is concentrated in the axes. These values increase depending on the habitat. Additionally, regardless of precipitation period and moment of the day, these modulation statistics show opposite behaviours overall, with one increasing while the other decreases. This indicates that modulation power is not spread across the MPS but is predominantly concentrated in either the low-pass region or on both axes of the MPS. Additionally, differences across habitats for **Starriness** remain less extreme and these variations may be explained by differences in geophony (as spectro-temporal modulations for biophony are more constrained). Nevertheless, Starriness may correlate with distance to the Equator as it shows an increasing tendency pointing in this direction, meaning that the closer to the Equator (habitats 8 and 9) the higher Starriness values overall.
iii. **Separability** remains high for all habitats except for the habitats closer to the Equator (8 and 9). This may be related to the latitudinal biodiversity gradient as this is one of the main factors that so drastically distinguishes 8 and 9 from the other habitats. Also, higher biodiversity implies more animal choruses than isolated sound sources, which should also decrease Separability values.
iv. Similarly to modulation depth, **Alpha** shows some of its highest values for habitats 4 and 7 (midday) and lowest for 8 and 9. However, these variations are harder to interpret in light of the findings from contrasts of biophony/geophony and isolated/choruses sound sources. These findings suggest that biophony has higher Alpha values than geophony without animal choruses changing this behaviour, which points toward higher values of Alpha for more biodiverse habitats but is not what is observed in figure 5. Therefore, distinction between open and closed habitats remains the only habitat characteristic that can be associated with variations in Alpha due to sound propagation effects (higher values for open habitats, lower for closed ones).

Up to this point, most of these interpretations are not conclusive and should be considered as potential candidates to try to explain and relate part of the variability observed in modulation statistics to different characteristics of the habitats. More analyses are definitely needed to unveil and expand further our knowledge on how different habitat characteristics may influence the modulation information conveyed by natural soundscapes.

## IV. DISCUSSION

### A. Modulation statistics of natural soundscapes

Thoret *et al*. (2020) and Apoux *et al*. (2023) found that relatively slow spectro-temporal modulation cues conveyed by natural soundscapes provide sufficient information for accurate classification of wild habitats, irrespective of moment of the day and season. However, these studies were limited to four habitats of a single temperate biome (a protected nature reserve) in California (USA). The current study was based on a much larger database of acoustic samples recorded in nine terrestrial habitats, marginally affected by human activity, over five continents which were recorded at four moments of the day and two precipitation periods. This study showed that natural soundscapes are generally low-pass in shape in the modulation domain, with most of their modulation power restricted to low temporal (<10-20 Hz) and spectral modulations (<0.5-1 cycles/kHz). These spectro-temporal modulation cues differ across habitats, and more specifically as a function of the habitat’s distance to the equatorial region, irrespective of moment of the day and precipitation period. In other words, natural soundscapes convey valuable information in the spectrotemporal modulation domain that could support detection of acoustic changes within and across habitats with potentially behavioural relevance for human and nonhuman animals. Amongst the six modulation statistics proposed by Singh and Theunissen (2003), Modulation depth and Starriness contributed most to the distinction between these natural habitats. Moreover, Starriness and Separability distinguished natural soundscapes recorded in equatorial habitats from those recorded at other latitudes in a relatively robust manner (that is, irrespective of moment of the day and precipitation period). Overall, this result is consistent with expected (global) variations in animal acoustic activity associated with the well-known latitudinal gradient of biodiversity (e.g., Gaston, 2000). This result therefore suggests that biophony –the collective sound produced by biological sound sources and its variations (biodiversity, i.e., abundance and species richness)– may be characterised by the presence of joint spectro-temporal modulations together with the lack of joint high spectral and fast temporal modulations for biological sounds in their natural context. Joint spectro-temporal modulations (low Separability) may point towards presence of frequency sweeps and/or diversity of distinct sound sources, or in other words high MPS complexity. High Starriness may point to the specific bio-mechanical and resulting spectro-temporal constraints of biotic sound sources in the generation of vocal signals such as bird songs, amphibians or mammal calls and insect stridulations/timbalations.

Other modulation statistics such as asymmetry and alpha made a relatively modest contribution to the distinction between natural habitats. Asymmetry was close to zero for all habitats. This suggests that modulation spectra of natural soundscapes –perhaps due to the combination of all kinds of animal vocalisations– have equal representation of down-sweeps and up-sweeps. The small contribution of alpha is more surprising given the large number of studies showing that animal vocalisations follow 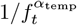 temporal-modulation power-law relationships (Attias and Schreiner, 1996; Jermyn *et al*., 2023; Singh and Theunissen, 2003) quite different from geophysical sounds (Attias and Schreiner, 1996; Geffen *et al*., 2011; Singh and Theunissen, 2003) or even rural soundscapes (De Coensel and Botteldooren, 2006; De Coensel *et al*., 2003). However Yang *et al*. (2015) discusses examples where environmental sounds do not seem to follow this relationship. This is unlikely due to the use of short-duration (1 sec) acoustic samples as similar 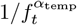 laws could be obtained for shorter samples of animal calls or speech sounds with the current MPS analysis. The smaller contribution of alpha is quite likely a consequence of combining animal vocalisations with geophysical sounds and applying specific sound-propagation processes in the same soundscape. If correct, this conclusion points to the importance of studying 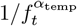 laws in contexts of high ecological validity.

Overall, these results extend the initial conclusions of previous studies (e.g., Attias and Schreiner, 1996; Singh and Theunissen, 2003) to complex and more ecologically valid acoustic scenes, but they also raise the difficulty of extending the results obtained with sounds extracted from their original context. The Hearbiodiv database is available for further ecoacoustic, computational, neuroscientific and behavioural studies upon request to the management.

### B. Implications for hearing sciences

A characterisation of modulation statistics in situations of high ecological validity should be useful for future psychoacoustical, neuroscientific and ethological investigations aiming at testing efficient-coding principles and/or identifying auditory cues and sensory mechanisms used by human and non-human animals for detecting presence, numerosity and variety of biological sound sources. The present results –and more specifically the important role played by i) the low spectro-temporal modulations and ii) lack of joint high spectral and temporal modulations– for classification of natural sounds are in line with pilot psychoacoustical results obtained in a group of human observers asked to detect presence of living being when listening to samples taken from the Hearbiodiv sound database (Miller-Viacava *et al*., 2023a,b)

Altogether, these findings impose specific constraints on the neural mechanisms involved in the monitoring of biological and geophysical sound sources in the near environment by the peripheral and central auditory system of human and non-human species (Depireux *et al*., 2001; Homma *et al*., 2020; Lorenzi *et al*., 2023) and more broadly, on the environment statistics that are used to build an internal model of the external world to make predictions of future acoustic events (e.g., Denham and Winkler, 2020; Skerritt-Davis and Elhilali, 2021a,b). These findings may also guide future work investigating the ability of human listeners to hear the global properties of natural scenes, in line with the pioneering work of Mc-Mullin *et al*. (2024) showing that humans consistently perceive openness (a structural property), transience (a constancy property), and navigability (a functional property) in real-world auditory scenes such as parks, forests and hiking trails.

### C. Implications for studies in soundscape ecology and ecoacoustics

The Hearbiodiv database and results from this study should also be useful for ecological studies searching for robust acoustic markers of biodiversity also called “ecoacoustic indices”. Recent studies demonstrated that most ecoacoustic indices developed as proxies of biodiversity for biodiversity monitoring are generally unsuitable for geographic comparisons (e.g., Giuliani *et al*., 2024) and fail at showing strong effects of latitude (Eldridge *et al*., 2018; Pan *et al*., 2024). This is partly because spectrotemporal overlap across species (birds, insects, amphibians) is expected to culminate in high-diversity environments such as tropical ones (Sueur *et al*., 2008), current ecoacoustic indices being too simplistic to alleviate this blind source separation problem. The decomposition of natural soundscapes in the spectro-temporal modulation domain –because of its high relevance to animal sounds and animal communication (Singh and Theunissen, 2003)– may provide an effective solution to this source separation problem (Grinfeder *et al*., 2024). The present findings should encourage ecoacousticians to apply MPS analysis on their databases and develop novel ecoacoustic indexes based on modulation statistics, in particular Modulation depth, Starriness and Separability.

## V. CONCLUSIONS

The present study estimated modulation statistics of natural soundscapes using a large database of natural scenes or soundscapes recorded in nine distinct terrestrial habitats. Altogether, the analyses showed that:

1. The modulation power spectra of these natural soundscapes were generally low-pass in shape, with most of their modulation power restricted to slow temporal modulations (<10-20 Hz) and low spectral modulations (<0.5-1 cycles/kHz).
2. The modulation statistics calculated from the modulation power spectra of these soundscapes enabled natural habitats to be distinguished in a robust manner.
3. Two modulation statistics –Modulation depth and Starriness– contributed most to the distinction between natural habitats. Moreover, Starriness and Separability distinguished the most biodiverse (i.e., equatorial) habitats from the remaining ones, consistent with the idea that biotic sounds impose specific constraints (i.e., they cannot show rapid temporal modulation and high spectral modulations simultaneously) on the spectro-temporal structure of natural soundscapes.
4. A modulation statistic often used to characterise natural sounds and scenes, the so-called “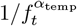 temporal-modulation power-law relationship”, proved to be poorly effective in its ability to distinguish habitats, illustrating the importance of characterising the regularities of natural sounds in their original context and using large, ecologically-valid databases.

This study demonstrates the value of using the “modulation theory” (Singh and Theunissen, 2003) to describe the statistical properties of natural soundscapes and its potential for studying ecological processes underlying these complex acoustic scenes. It corroborates the idea that terrestrial biomes have a unique “acoustic signature” in the spectro-temporal domain that may be unveiled by adequate modulation analysis.

## ACKNOWLEDGMENTS

This research was supported by FrontCog ANR-17-EURE-0017, ANR-20-CE28-0011 and ANR-20-CE28- 0011 Hearbiodiv. Miller-Viacava was supported by a doctoral grant from ED540 ENS-PSL. The work conducted in Haut Jura was supported by the Parc Naturel Régional du Haut-Jura, which received funding from the Région Bourgogne-Franche-Comté, the Région Auvergne-Rhône-Alpes, and the DREAL Bourgogne-Franche-Comté. The work conducted in Nouragues was supported by Labex CEBA which received funding from Investissements d’Avenir ANR-10-LABX-25-01. Work in La Belgique was done thanks to funding provided by the Flemish Government to the Antwerp Zoo Centre for Research and Conservation.

The authors wish to thank Richard McWalter and Elie Grinfeder for their valuable input on this work. Additionaly, to Anne C. Axel (Twin Lakes), Diego Llusia (Chapada dos Veadeiros) and Yvonne F. Phillips (Woondum) for sharing natural soundscapes recordings, and to Bernie Krause (Wild sanctuary, Sonoma, USA) for sharing one river sound.

Concerning the soundscapes recordings the authors thank: (i) for Haut Jura, Sylvain Haupert, Frédéric Sèbe, Marie-Pierre Reynet and Julien Barlet; (ii) for Ironwood, François-Michel Le Tourneau and Ruth Gosset; (iii) for Yanbaru, Samuel Ross, and special thanks to the landowners, museums, local governments and schools that host the OKEON Churamori project, to the people of Okinawa; (iv) for Nouragues, Philippe Gaucher, Olivier Claessens, Elodie Courtois, Nina Marchand and Elodie Schloesing; (v) for La Belgique, Jacques Keumo Kuenbou and Lena Kuperus.

The findings and conclusions in this article are those of the authors and do not necessarily represent the views of any government agencies. Any use of trade, product, or firm names is for descriptive purposes only and does not imply and endorsement by the U.S. government

## DATA AVAILABILITY

The sounds in the Hearbiodiv database created for this study are available upon request to the management. All data discussed on this study, and codes created for proccessing, are available upon request to the corresponding author.

## APPENDIX A EFFECTS OF SOUND SAMPLE DURATION

To assess possible effects of sound sample duration on modulation statistics computed for the Hearbiodiv database, the same sound sample selection as explained in section IIA3 was conducted but instead of selecting

4 samples of 1-sec long each, only 1 sample of 1-min long was selected for each moment of the day. This means that the total number of samples analysed with longer duration is 2880 (11520/4). Modulation statistics for longer samples are presented on figure 9. It is clear that the range of values for each modulation statistic is not exactly the same compared to results obtained for 1-sec long samples shown before on figure 5. Still, what is most relevant here are the differences observed across habitats, as this relates to the possibility of using modulation statistics to distinguish across habitats. To better compare the trends across habitats, modulation statistics obtained for both 1-sec and 1-min long samples were normalised by the maximum value of each statistic which is presented in figure 10. These results show that the trends across habitats remain fairly similar for both sample duration, especially for Modulation depth and Starriness which were found to contribute the most to habitat classification. The most noticeable differences are observed in Separability for habitats closer to the Equator (8 and 9) which become highly separable, Lowpass coefficient which keeps the same distribution across habitats but with much higher differences, with habitats 2, 4, 5 and 7 showing much lower values (i.e., modulation power less concentrated in the low-pass region), and Asymmetry which differences depend on precipitation period.

**FIG. 9.**
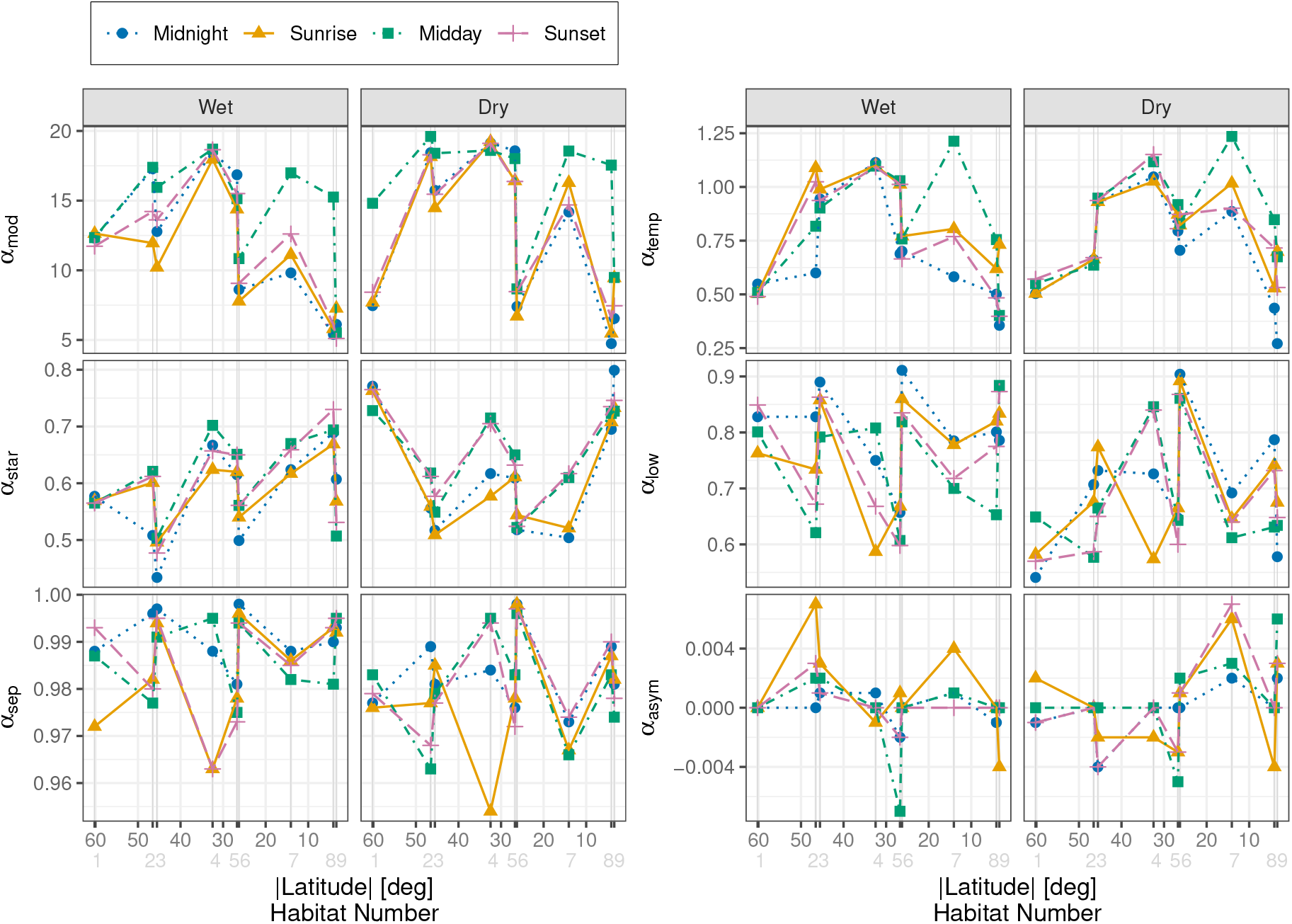
Same as figure 5 but for 2880 1-min long sound samples.

**FIG. 10.**
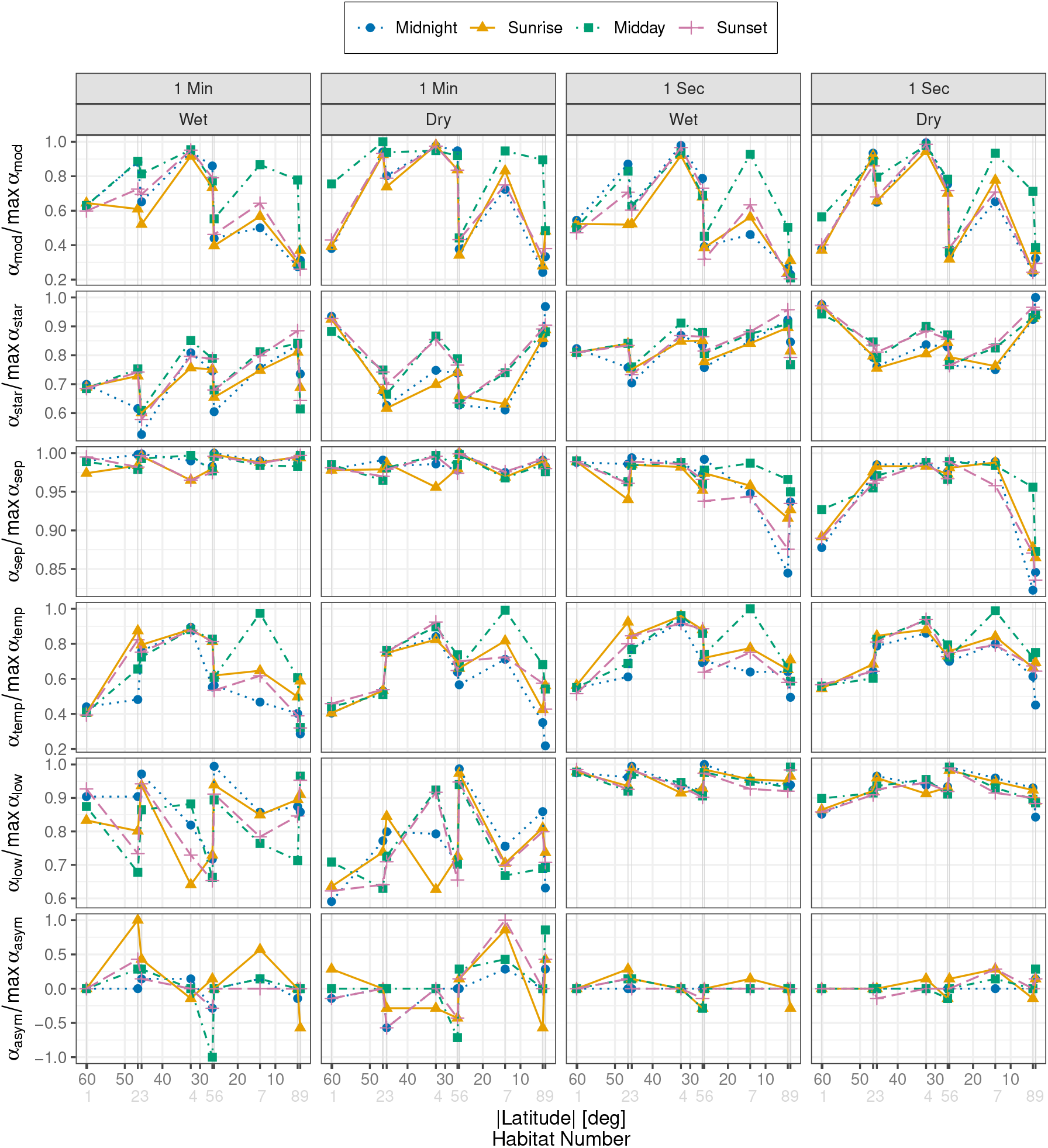
Same as figures 9 (two columns left) and 5 (two columns right) but each plot was normalised by its maximum value to allow better comparison across sound sample duration.

## APPENDIX B ART ANOVA AND STATISTICAL ANALYSIS

### About normal distributions of data

Although Shapiro-Wilk test tends to reject normality for larger sample sizes (as small deviations from normality would have a big impact on significance assesement), it remains a simple way to ilustrate and quantify normality. It shows that from all 72 combinations (9 habitats x 2 precipitation periods x 4 moments of the day), the percentage of the ones with significant p-values (meaning non normally distributed) are generally high overall: Modulation depth: 90.3%, Starriness: 79.2%, Separability: 100%, Alpha: 88.9%, Low-pass coefficient: 88.9%, Asymmetry: 91.7%. These results were further confirmed with QQ-plots of the 72 combinations for each modulation statistic. This is consistent with what can be seen in figure 4 with the density distribution estimations of the modulation statistics values for all samples in the Hearbiodiv database.

### About ART ANOVA

Concerning the analysis for studying the differences observed across habitats, moments of the day, and precipitation periods, it was found that the distributions of modulation statistics violated almost all ANOVA assumptions: (i) the data is not completely independent as some sound samples come from the same day, (ii) studied with QQ-plots, residuals were not normally distributed for Separability, Asymmetry and Alpha, and to a lesser extent for Modulation Depth and Starriness, (iii) as expected from the distribution plots shown on figure 4, homogeneity of the variances was violated by all modulation statistics with Leverne’s test showing significant *p*-values (< 2.2 × 10^−16^) for all of them. Therefore, a non-parametric method was needed that could manage tree main factors (habitat, moment of the day and rainfall period). Conducting an ANOVA on ranked transformed data is one of the many possible approaches when an alternative to parametric tests is needed. This approach has shown to be successful in the past although there are still ongoing debates on its feasibility when experimental designs involve more complex combination with higher factor numbers and/or modalities of each factor (e.g., Conover and Iman, 1981; Leys and Schumann, 2010; Sawilowsky, 1990; Sawilowsky *et al*., 1989; Tsandilas and Casiez, 2025, and references therein). Still, this statistical approach remains the most straightforward and easy to interpret in many different fields method, which was considered an important asset due to the interdisciplinary nature of this study.

As some researchers suggest that regular ANOVA may be more robust to slight violations of its assumptions compared to the potential problems of using ART ANOVA with large number of factors, interactions were checked also with a classical parametric three-way ANOVA and two-way ANOVA (the subsequent two-way ANOVAs were calculated with errors associated only to the groups being studied in each case and not the pooled error from the full factorial design due to the presence of heterogeneity of variance in the data, following the recommendations from Keppel and Wickens (2004). The same was done for results of the two-way interactions using the ART ANOVA in the Results section). These interactions were also found to be significant as shown on tables V, VI and VII.

**TABLE V.** Significant three-way interactions (Habitat:Moment:Period) resulting from all six ANOVAs, one for each modulation statistic.

| Mod Stat | DFn | DFd | F | p | $p \leq .05$ |
| --- | --- | --- | --- | --- | --- |
| $\alpha_{\text{mod}}$ | 24 | 11448 | 13.729 | 1.39e-54 | * |
| $\alpha_{\text{star}}$ | 24 | 11448 | 10.561 | 1.08e-39 | * |
| $\alpha_{\text{sep}}$ | 24 | 11448 | 16.899 | 1.24e-69 | * |
| $\alpha_{\text{temp}}$ | 24 | 11448 | 10.835 | 5.67e-41 | * |
| $\alpha_{\text{low}}$ | 24 | 11448 | 6.454 | 5.94e-21 | * |
| $\alpha_{\text{asym}}$ | 24 | 11448 | 3.405 | 3.5e-08 | * |

**TABLE VI.** Two-way interactions (Habitat:Moment) for each precipitation period resulting from six follow up ANOVAs, one for each modulation statistic. Significance level of 0.008 after Bonferroni correction by the number of two-way interaction computed (6).

| Mod Stat | Period | DFn | DFd | F | p | $p \leq .008$ |
| --- | --- | --- | --- | --- | --- | --- |
| $\alpha_{\text{mod}}$ | Wet | 24 | 5724 | 27.9 | 1.86e-118 | * |
| $\alpha_{\text{mod}}$ | Dry | 24 | 5724 | 25.8 | 4.24e-109 | * |
| $\alpha_{\text{star}}$ | Wet | 24 | 5724 | 18.0 | 1.66e-73 | * |
| $\alpha_{\text{star}}$ | Dry | 24 | 5724 | 31.8 | 5.92e-136 | * |
| $\alpha_{\text{sep}}$ | Wet | 24 | 5724 | 42.1 | 2.60e-181 | * |
| $\alpha_{\text{sep}}$ | Dry | 24 | 5724 | 63.0 | 5.25e-269 | * |
| $\alpha_{\text{temp}}$ | Wet | 24 | 5724 | 22.6 | 2.31e-94 | * |
| $\alpha_{\text{temp}}$ | Dry | 24 | 5724 | 12.5 | 3.35e-48 | * |
| $\alpha_{\text{low}}$ | Wet | 24 | 5724 | 12.8 | 1.97e-49 | * |
| $\alpha_{\text{low}}$ | Dry | 24 | 5724 | 21.4 | 7.26e-89 | * |
| $\alpha_{\text{asym}}$ | Wet | 24 | 5724 | 3.87 | 5.95e-10 | * |
| $\alpha_{\text{asym}}$ | Dry | 24 | 5724 | 6.47 | 7.34e-21 | * |

**TABLE VII.**
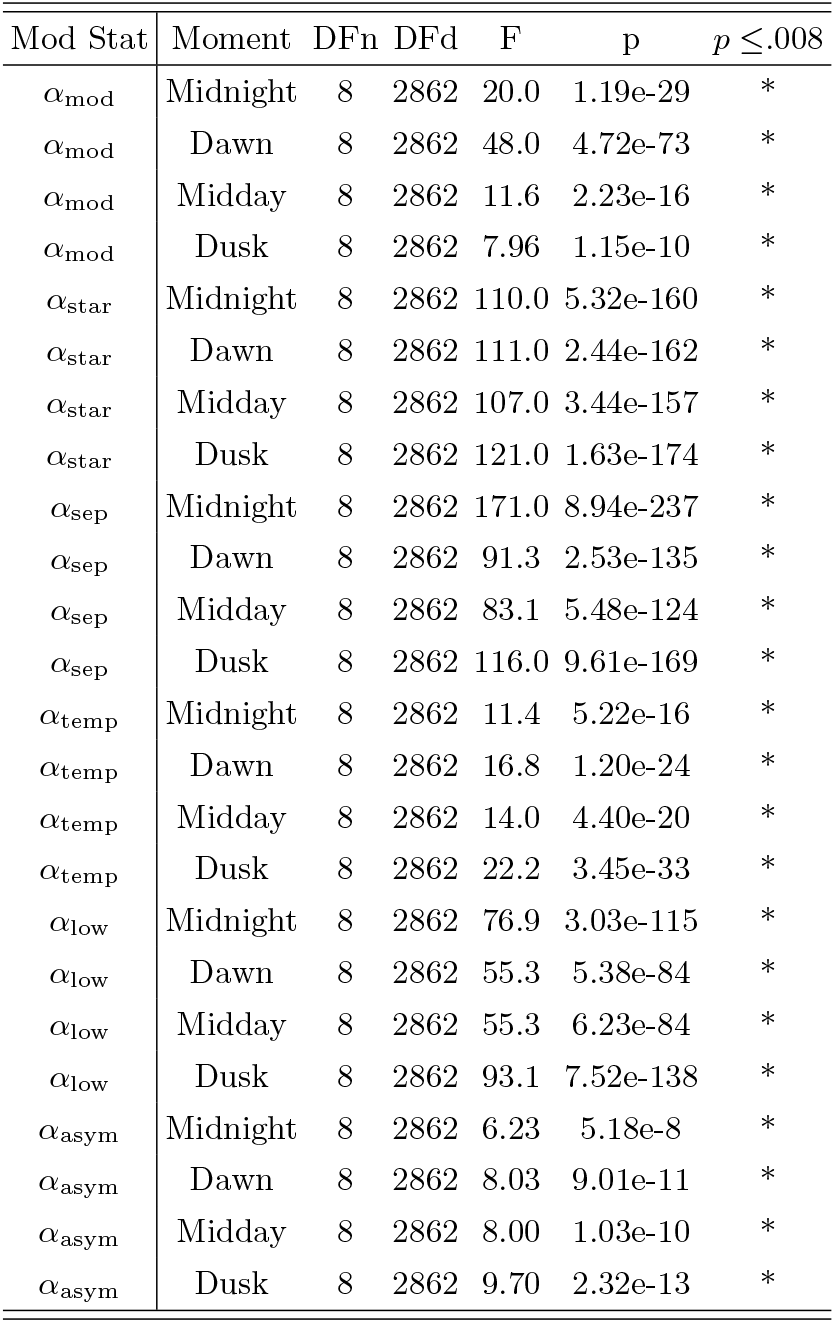
Two-way interactions (Habitat:Period) for each moment of the day resulting from six follow up ANOVAs, one for each modulation statistic. Significance level of 0.008 after Bonferroni correction by the number of two-way interaction computed (6).

| Mod Stat | Moment | DFn | DFd | F | p | $p \leq .008$ |
| --- | --- | --- | --- | --- | --- | --- |
| $\alpha_{\text{mod}}$ | Midnight | 8 | 2862 | 20.0 | 1.19e-29 | * |
| $\alpha_{\text{mod}}$ | Dawn | 8 | 2862 | 48.0 | 4.72e-73 | * |
| $\alpha_{\text{mod}}$ | Midday | 8 | 2862 | 11.6 | 2.23e-16 | * |
| $\alpha_{\text{mod}}$ | Dusk | 8 | 2862 | 7.96 | 1.15e-10 | * |
| $\alpha_{\text{star}}$ | Midnight | 8 | 2862 | 110.0 | 5.32e-160 | * |
| $\alpha_{\text{star}}$ | Dawn | 8 | 2862 | 111.0 | 2.44e-162 | * |
| $\alpha_{\text{star}}$ | Midday | 8 | 2862 | 107.0 | 3.44e-157 | * |
| $\alpha_{\text{star}}$ | Dusk | 8 | 2862 | 121.0 | 1.63e-174 | * |
| $\alpha_{\text{sep}}$ | Midnight | 8 | 2862 | 171.0 | 8.94e-237 | * |
| $\alpha_{\text{sep}}$ | Dawn | 8 | 2862 | 91.3 | 2.53e-135 | * |
| $\alpha_{\text{sep}}$ | Midday | 8 | 2862 | 83.1 | 5.48e-124 | * |
| $\alpha_{\text{sep}}$ | Dusk | 8 | 2862 | 116.0 | 9.61e-169 | * |
| $\alpha_{\text{temp}}$ | Midnight | 8 | 2862 | 11.4 | 5.22e-16 | * |
| $\alpha_{\text{temp}}$ | Dawn | 8 | 2862 | 16.8 | 1.20e-24 | * |
| $\alpha_{\text{temp}}$ | Midday | 8 | 2862 | 14.0 | 4.40e-20 | * |
| $\alpha_{\text{temp}}$ | Dusk | 8 | 2862 | 22.2 | 3.45e-33 | * |
| $\alpha_{\text{low}}$ | Midnight | 8 | 2862 | 76.9 | 3.03e-115 | * |
| $\alpha_{\text{low}}$ | Dawn | 8 | 2862 | 55.3 | 5.38e-84 | * |
| $\alpha_{\text{low}}$ | Midday | 8 | 2862 | 55.3 | 6.23e-84 | * |
| $\alpha_{\text{low}}$ | Dusk | 8 | 2862 | 93.1 | 7.52e-138 | * |
| $\alpha_{\text{asym}}$ | Midnight | 8 | 2862 | 6.23 | 5.18e-8 | * |
| $\alpha_{\text{asym}}$ | Dawn | 8 | 2862 | 8.03 | 9.01e-11 | * |
| $\alpha_{\text{asym}}$ | Midday | 8 | 2862 | 8.00 | 1.03e-10 | * |
| $\alpha_{\text{asym}}$ | Dusk | 8 | 2862 | 9.70 | 2.32e-13 | * |

In any case, the consistency between results obtained with ART ANOVA and subsequent machine learning classifier supports that the results found in this study concerning habitat classification are relevant and not due to potential statistical errors. Still, further reasearch could be done to explore the influence of habitat, moment of the day, and precipitation period on each modulation statistics with more tailored statistical methods suited to the high complexity of the data obtained in this study.

## BIBLIOGRAPHY

Amézquita, A., and Hödl, W. (2004). “How, when, and where to perform visual displays: The case of the Amazonian frog Hyla parviceps,” Herpetologica 60(4), 420–429, mu doi: 10.1655/02-51.

Apoux, F., Miller-Viacava, N., Ferriére, R., Dai, H., Krause, B., Sueur, J., and Lorenzi, C. (2023). “Auditory discrimination of natural soundscapes,” The Journal of the Acoustical Society of America 153(5), 2706, mu doi: 10.1121/10.0017972.

Ardoint, M., Lorenzi, C., Pressnitzer, D., and Gorea, A. (2008). “Investigation of perceptual constancy in the temporal-envelope domain,” The Journal of the Acoustical Society of America 123(3), 1591–1601, mu doi: 10.1121/1.2836782.

Attias, H., and Schreiner, C. (1996). “Temporal Low-Order Statistics of Natural Sounds,” in Advances in Neural Information Processing Systems, edited by M. C. Mozer, M. Jordan, and T. Petsche, MIT Press, Vol. 9.

Attneave, F. (1954). “Some informational aspects of visual perception.,” Psychological Review 61(3), 183–193, mu doi: 10.1037/h0054663.

Aubin, T., and Bremond, J.-C. (1983). “The Process of Species-specific Song Recognition in the Skylark Alauda arvensis. An Experimental Study by Means of Synthesis,” Zeitschrift für Tierpsychologie 61(2), 141–152, mu doi: 10.1111/j.1439-0310.1983.tb01334.x.

Barlow, H. B. (1961). Possible principles underlying the transformation of sensory messages. Sensory communication: contributions to the Symposium on principles of sensory communication, 1(01) (the MIT press, Cambridge (Mass.)).

Bedoya, C., Isaza, C., Daza, J. M., and López, J. D. (2017). “Automatic identification of rainfall in acoustic recordings,” Ecological Indicators 75, 95–100, mu doi: 10.1016/j.ecolind.2016.12.018.

Brown, J. H., Gillooly, J. F., Allen, A. P., Savage, V. M., and West, G. B. (2004). “Toward a metabolic theory of ecology,” Ecology 85(7), 1771–1789, mu doi: 10.1890/03-9000.

Capranica, R. R., Rose, G. J., and Brenowitz, E. A. (1985). “Time Resolution in the Auditory Systems of Anurans,” in Time Resolution in Auditory Systems, edited by A. Michelsen (Springer Berlin Heidelberg, Berlin, Heidelberg), pp. 58–73, mu doi: 10.1007/978-3-642-70622-6_4.

Catchpole, C. K., and Slater, P. J. B. (2008). Bird Song: Biological Themes and Variations, 2 ed. (Cambridge University Press).

Chen, Y.-F., Luo, Y., Mammides, C., Cao, K.-F., Zhu, S., and Goodale, E. (2021). “The relationship between acoustic indices, elevation, and vegetation, in a forest plot network of southern China,” Ecological Indicators 129, 107942, mu doi: 10.1016/j.ecolind.2021.107942.

Chi, T., Gao, Y., Guyton, M. C., Ru, P., and Shamma, S. (1999). “Spectro-temporal modulation transfer functions and speech intelligibility,” The Journal of the Acoustical Society of America 106(5), 2719–2732, mu doi: 10.1121/1.428100.

Conover, W. J., and Iman, R. L. (1981). “Rank Transformations as a Bridge between Parametric and Nonparametric Statistics,” The American Statistician 35(3), 124–129, mu doi: 10.1080/00031305.1981.10479327.

De Baudouin, A., Couprie, P., Michaud, F., Haupert, S., and Sueur, J. (2024). “Similarity visualization of soundscapes in ecology and music,” Frontiers in Ecology and Evolution 12, 1334776, mu doi: 10.3389/fevo.2024.1334776.

De Coensel, B., and Botteldooren, D. (2006). “The quiet rural soundscape and how to characterize it,” Acta Acustica united with Acustica 92(6), 887–897.

De Coensel, B., Botteldooren, D., and De Muer, T. (2003). “1/f Noise in Rural and Urban Soundscapes,” Acta Acustica united with Acustica 89(2), 287–295.

Denham, S. L., and Winkler, I. (2020). “Predictive coding in auditory perception: Challenges and unresolved questions,” European Journal of Neuroscience 51(5), 1151–1160, mu doi: 10.1111/ejn.13802.

Depireux, D. A., Simon, J. Z., Klein, D. J., and Shamma, S. A. (2001). “Spectro-Temporal Response Field Characterization With Dynamic Ripples in Ferret Primary Auditory Cortex,” Journal of Neurophysiology 85(3), 1220–1234, mu doi: 10.1152/jn.2001.85.3.1220.

Diepstraten, J., and Willie, J. (2021). “Assessing the structure and drivers of biological sounds along a disturbance gradient,” Global Ecology and Conservation 31, e01819, mu doi: 10.1016/j.gecco.2021.e01819.

Divyapriya, C., and Pramod, P. (2019). “Ornithophony in the soundscape of Anaikatty Hills, Coimbatore, Tamil Nadu, India,” Journal of Threatened Taxa 11(12), 14471–14483, mu doi: 10.11609/jott.4948.11.12.14471-14483.

Dooling, R. J., and Popper, A. N. (2000). “Hearing in Birds and Reptiles: An Overview,” in Comparative Hearing: Birds and Reptiles, edited by R. R. Fay, A. N. Popper, R. J. Dooling, R. R. Fay, and A. N. Popper, 13 (Springer New York, New York, NY), pp. 1–12, mu doi: 10.1007/978-1-4612-1182-2_1.

Droste, R. (2022). “sunriseset(lat, lng, utcoff, date, plot)” muhttps://www.mathworks.com/matlabcentral/fileexchange/62180-sunriseset-lat-lng-utcoff-date-plot, [MATLAB Central File Exchange. Retrieved January 3, 2022].

Drullman, R. (1995). “Temporal envelope and fine structure cues for speech intelligibility,” The Journal of the Acoustical Society of America 97(1), 585–592, mu doi: 10.1121/1.413112.

Eggermont, J. J. (2002). “Temporal Modulation Transfer Functions in Cat Primary Auditory Cortex: Separating Stimulus Effects From Neural Mechanisms,” Journal of Neurophysiology 87(1), 305–321, mu doi: 10.1152/jn.00490.2001.

Eldridge, A., Guyot, P., Moscoso, P., Johnston, A., Eyre-Walker, Y., and Peck, M. (2018). “Sounding out ecoacoustic metrics: Avian species richness is predicted by acoustic indices in temperate but not tropical habitats,” Ecological Indicators 95, 939–952, mu doi: 10.1016/j.ecolind.2018.06.012.

Elkin, L. A., Kay, M., Higgins, J. J., and Wobbrock, J. O. (2021). “An Aligned Rank Transform Procedure for Multifactor Contrast Tests,” in The 34th Annual ACM Symposium on User Interface Software and Technology, ACM, Virtual Event USA, pp. 754– 768, mu doi: 10.1145/3472749.3474784.

Elliott, T. M., and Theunissen, F. E. (2009). “The Modulation Transfer Function for Speech Intelligibility,” PLoS Computational Biology 5(3), e1000302, mu doi: 10.1371/journal.pcbi.1000302.

Escabí, M. A., Miller, L. M., Read, H. L., and Schreiner, C. E. (2003). “Naturalistic Auditory Contrast Improves Spectrotemporal Coding in the Cat Inferior Colliculus,” The Journal of Neuroscience 23(37), 11489–11504, mu doi: 10.1523/JNEUROSCI.23-37-11489.2003.

Farina, A., and Gage, S. H. (2017). Ecoacoustics: The Ecological Role of Sounds, 1st ed ed. (John Wiley & Sons, Inc, Hoboken, NJ).

Farina, A., Mullet, T. C., Bazarbayeva, T. A., Tazhibayeva, T., Bulatova, D., and Li, P. (2021). “Perspectives on the Ecological Role of Geophysical Sounds,” Frontiers in Ecology and Evolution 9, 748398, mu doi: 10.3389/fevo.2021.748398.

Fay, R. R., and Simmons, A. M. (1999). “The Sense of Hearing in Fishes and Amphibians,” in Comparative Hearing: Fish and Amphibians, edited by R. R. Fay, A. N. Popper, R. R. Fay, and A. N. Popper, 11 (Springer New York, New York, NY), pp. 269– 318, mu doi: 10.1007/978-1-4612-0533-3_7.

Field, D. J. (1987). “Relations between the statistics of natural images and the response properties of cortical cells,” Journal of the Optical Society of America A 4(12), 2379, mu doi: 10.1364/JOSAA.4.002379.

Field, D. J. (1994). “What Is the Goal of Sensory Coding?,” Neural Computation 6(4), 559–601, mu doi: 10.1162/neco.1994.6.4.559.

Fonseca, P. J. (2014). “Cicada Acoustic Communication,” in Insect Hearing and Acoustic Communication, edited by B. Hedwig, 1 (Springer Berlin Heidelberg, Berlin, Heidelberg), pp. 101–121, mu doi: 10.1007/978-3-642-40462-7_7.

Forrest, T. G. (1994). “From Sender to Receiver: Propagation and Environmental Effects on Acoustic Signals,” American Zoologist 34(6), 644–654, mu doi: 10.1093/icb/34.6.644.

Freitas, B., D’Amelio, P. B., Milá, B., Thébaud, C., and Janicke, T. (2024). “Meta-analysis of the acoustic adaptation hypothesis reveals no support for the effect of vegetation structure on acoustic signalling across terrestrial vertebrates,” Biological Reviews brv.13163, mu doi: 10.1111/brv.13163.

Fu, Q.-J., Galvin, J. J., and Wang, X. (2001). “Recognition of time-distorted sentences by normal-hearing and cochlear-implant listeners,” The Journal of the Acoustical Society of America 109(1), 379–384, mu doi: 10.1121/1.1327578.

Gage, S. H., and Axel, A. C. (2014). “Visualization of temporal change in soundscape power of a Michigan lake habitat over a 4-year period,” Ecological Informatics 21, 100–109, mu doi: 10.1016/j.ecoinf.2013.11.004.

Garcia-Lazaro, J., Ahmed, B., and Schnupp, J. (2006). “Tuning to Natural Stimulus Dynamics in Primary Auditory Cortex,” Current Biology 16(3), 264–271, mu doi: 10.1016/j.cub.2005.12.013.

Garcia-Lazaro, J. A., Ahmed, B., and Schnupp, J. W. H. (2011). “Emergence of Tuning to Natural Stimulus Statistics along the Central Auditory Pathway,” PLoS ONE 6(8), e22584, mu doi: 10.1371/journal.pone.0022584.

Gasc, A., Anso, J., Sueur, J., Jourdan, H., and Desutter-Grandcolas, L. (2018a). “Cricket calling communities as an indicator of the invasive ant Wasmannia auropunctata in an insular biodiversity hotspot,” Biological Invasions 20(5), 1099–1111, mu doi: 10.1007/s10530-017-1612-0.

Gasc, A., Gottesman, B. L., Francomano, D., Jung, J., Durham, M., Mateljak, J., and Pijanowski, B. C. (2018b). “Soundscapes reveal disturbance impacts: Biophonic response to wildfire in the Sonoran Desert Sky Islands,” Landscape Ecology 33(8), 1399– 1415, mu doi: 10.1007/s10980-018-0675-3.

Gaston, K. J. (2000). “Global patterns in biodiversity,” Nature 405(6783), 220–227, mu doi: 10.1038/35012228.

Gaston, K. J., and Blackburn, T. M., eds. (2000). Pattern and Process in Macroecology, 1 ed. (Wiley).

Geffen, M. N., Gervain, J., Werker, J. F., and Magnasco, M. O. (2011). “Auditory Perception of Self-Similarity in Water Sounds,” Frontiers in Integrative Neuroscience 5, mu doi: 10.3389/fnint.2011.00015.

Gerhardt, H. C., and Bee, M. A. (2006). “Recognition and Localization of Acoustic Signals,” in Hearing and Sound Communication in Amphibians, edited by P. M. Narins, A. S. Feng, R. R. Fay, and A. N. Popper, 28 (Springer New York), pp. 113–146, mu doi: 10.1007/978-0-387-47796-1_5.

Gil, D., and Llusia, D. (2020). “The Bird Dawn Chorus Revisited,” in Coding Strategies in Vertebrate Acoustic Communication, edited by T. Aubin and N. Mathevon, 7 (Springer International Publishing, Cham), pp. 45–90, mu doi: 10.1007/978-3-030-39200-0_3.

Giuliani, M., Mirante, D., Abbondanza, E., and Santini, L. (2024). “Acoustic indices fail to represent different facets of biodiversity,” Ecological Indicators 166, 112451, mu doi: 10.1016/j.ecolind.2024.112451.

Grinfeder, E., Haupert, S., Ducrettet, M., Barlet, J., Reynet, M.P., Sébe, F., and Sueur, J. (2022a). “Soundscape dynamics of a cold protected forest: Dominance of aircraft noise,” Landscape Ecology 37(2), 567–582, mu doi: 10.1007/s10980-021-01360-1.

Grinfeder, E., Lorenzi, C., Haupert, S., and Sueur, J. (2022b). “What Do We Mean by “Soundscape”? A Functional Description,” Frontiers in Ecology and Evolution 10, 894232, mu doi: 10.3389/fevo.2022.894232.

Grinfeder, E., Lorenzi, C., Teytaut, Y., Haupert, S., and Sueur, J. (2024). “Introducing Evascape, a Model-Based Soundscape Assembler: Impact of Background Sounds on Biodiversity Monitoring with Ecoacoustic Indices,” [preprint] 10.2139/ssrn.5045662.

He, F., Stevenson, I. H., and Escabí, M. A. (2023). “Two stages of bandwidth scaling drives efficient neural coding of natural sounds,” PLOS Computational Biology 19(2), e1010862, mu doi: 10.1371/journal.pcbi.1010862.

Holleman, G. A., Hooge, I. T. C., Kemner, C., and Hessels, R. S. (2020). “The ‘Real-World Approach’ and Its Problems: A Critique of the Term Ecological Validity,” Frontiers in Psychology 11, 721, mu doi: 10.3389/fpsyg.2020.00721.

Homma, N. Y., Hullett, P. W., Atencio, C. A., and Schreiner, C. E. (2020). “Auditory Cortical Plasticity Dependent on Environmental Noise Statistics,” Cell Reports 30(13), 4445–4458.e5, mu doi: 10.1016/j.celrep.2020.03.014.

Hsu, A., Woolley, S. M. N., Fremouw, T. E., and Theunissen, F. E. (2004). “Modulation Power and Phase Spectrum of Natural Sounds Enhance Neural Encoding Performed by Single Auditory Neurons,” The Journal of Neuroscience 24(41), 9201–9211, mu doi: 10.1523/JNEUROSCI.2449-04.2004.

Jermyn, A. S., Stevenson, D. J., and Levitin, D. J. (2023). “1/f laws found in non-human music,” Scientific Reports 13(1), 1324, mu doi: 10.1038/s41598-023-28444-z.

Joris, P. X., Schreiner, C. E., and Rees, A. (2004). “Neural Processing of Amplitude-Modulated Sounds,” Physiological Reviews 84(2), 541–577, mu doi: 10.1152/physrev.00029.2003.

Keppel, G., and Wickens, T. D. (2004). Design and Analysis: A Researcher’s Handbook, 4rd Ed (Pearson Prentice-Hall, Upper Saddle River, NJ).

Kohlrausch, A., Fassel, R., and Dau, T. (2000). “The influence of carrier level and frequency on modulation and beat-detection thresholds for sinusoidal carriers,” The Journal of the Acoustical Society of America 108(2), 723–734, mu doi: 10.1121/1.429605.

Krause, B. (1987). “Bioacoustics, habitat ambience in ecological balance,” Whole Earth Review 57(472), 14–18.

Lack, D. (1950). “Breeding seasons in the Galapagos.,” Ibis 92(2), 268–278, mu doi: 10.1111/j.1474-919X.1950.tb01751.x.

Lengagne, T., and Slater, P. J. B. (2002). “The effects of rain on acoustic communication: Tawny owls have good reason for calling less in wet weather,” Proceedings of the Royal Society of London. Series B: Biological Sciences 269(1505), 2121–2125, mu doi: 10.1098/rspb.2002.2115.

Lesica, N. A., and Grothe, B. (2008). “Efficient Temporal Processing of Naturalistic Sounds,” PLoS ONE 3(2), e1655, mu doi: 10.1371/journal.pone.0001655.

Lewicki, M. S. (2002). “Efficient coding of natural sounds,” Nature Neuroscience 5(4), 356–363, mu doi: 10.1038/nn831.

Leys, C., and Schumann, S. (2010). “A nonparametric method to analyze interactions: The adjusted rank transform test,” Journal of Experimental Social Psychology 46(4), 684–688, mu doi: 10.1016/j.jesp.2010.02.007.

Lomolino, M. V., Riddle, B. R., and Whittaker, R. J. (2018). Biogeography: Biological Diversity across Space and Time, fifth edition ed. (Sinauer Associates, Inc., Publishers, Sunderland, Massachusetts, U.S.A).

Long, G. R. (1994). “Psychoacoustics,” in Comparative Hearing: Mammals, edited by R. R. Fay, A. N. Popper, R. R. Fay, and A. N. Popper, 4 (Springer New York,New York, NY), pp. 18– 56, mu doi: 10.1007/978-1-4612-2700-7_2.

Lorenzi, C., Apoux, F., Grinfeder, E., Krause, B., Miller-Viacava, N., and Sueur, J. (2023). “Human Auditory Ecology: Extending Hearing Research to the Perception of Natural Soundscapes by Humans in Rapidly Changing Environments,” Trends in Hearing 27, 23312165231212032, mu doi: 10.1177/23312165231212032.

McDermott, J. H., Oxenham, A. J., and Simoncelli, E. P. (2009). “Sound texture synthesis via filter statistics,” in 2009 IEEE Workshop on Applications of Signal Processing to Audio and Acoustics, IEEE, New Paltz, NY, USA, pp. 297–300, mu doi: 10.1109/ASPAA.2009.5346467.

McDermott, J. H., and Simoncelli, E. P. (2011). “Sound Texture Perception via Statistics of the Auditory Periphery: Evidence from Sound Synthesis,” Neuron 71(5), 926–940, mu doi: 10.1016/j.neuron.2011.06.032.

McMullin, M. A., Kumar, R., Higgins, N. C., Gygi, B., Elhilali, M., and Snyder, J. S. (2024). “Preliminary Evidence for Global Properties in Human Listeners During Natural Auditory Scene Perception,” Open Mind 8, 333–365, mu doi: 10.1162/opmi_a_00131.

McWalter, R., and Dau, T. (2017). “Cascaded Amplitude Modulations in Sound Texture Perception,” Frontiers in Neuroscience 11, 485, mu doi: 10.3389/fnins.2017.00485.

Meeus, J. (1991). Astronomical Algorithms (Willmann-Bell, Richmond (Va.)).

Michelsen, A., and Larsen, O. N. (1983). “Strategies for Acoustic Communication in Complex Environments,” in Neuroethology and Behavioral Physiology, edited by F. Huber and H. Markl (Springer Berlin Heidelberg, Berlin, Heidelberg), pp. 321–331, mu doi: 10.1007/978-3-642-69271-0_23.

Miller-Viacava, N., Axel, A. C., Ferriere, R., Friedman, N. R., Le Tourneau, F.-M., Llusia, D., Mullet, T. C., Phillips, Y. F., Willie, J., Sueur, J., and Lorenzi, C. (2023a). “Acoustic cues for biological sound-source perception by human listeners,” [Poster presentation] Basic Auditory Science. London, United Kingdom.

Miller-Viacava, N., Axel, A. C., Ferriere, R., Friedman, N. R., Le Tourneau, F.-M., Llusia, D., Mullet, T. C., Phillips, Y. F., Willie, J., Sueur, J., and Lorenzi, C. (2023b). “Categorisation of biological sounds in pristine soundscapes: A psychophysical investigation based on an ecologically valid database,” [Poster presentation] 46th ARO MidWinter Meeting. Orlando, Florida, United States.

Miller-Viacava, N., Lazard, D., Delmas, T., Krause, B., Apoux, F., and Lorenzi, C. (2024). “Sensorineural hearing loss alters auditory discrimination of natural soundscapes,” International Journal of Audiology 63(10), 809–818, mu doi: 10.1080/14992027.2023.2272559.

Moore, B. C. J. (2013). An Introduction to the Psychology of Hearing, 6. aufl ed. (Brill, Leiden, Boston).

Morton, E. S. (1975). “Ecological Sources of Selection on Avian Sounds,” The American Naturalist 109(965), 17–34, mu doi: 10.1086/282971.

Mouterde, S. C., Theunissen, F. E., Elie, J. E., Vignal, C., and Mathevon, N. (2014). “Acoustic Communication and Sound Degradation: How Do the Individual Signatures of Male and Female Zebra Finch Calls Transmit over Distance?,” PLoS ONE 9(7), e102842, mu doi: 10.1371/journal.pone.0102842.

Mullet, T. C., Farina, A., Morton, J. M., and Wilhelm, S. R. (2024). “Seasonal sonic patterns reveal phenological phases (sonophases) associated with climate change in subarctic Alaska,” Frontiers in Ecology and Evolution 12, 1345558, mu doi: 10.3389/fevo.2024.1345558.

Mullet, T. C., Gage, S. H., Morton, J. M., and Huettmann, F. (2016). “Temporal and spatial variation of a winter soundscape in south-central Alaska,” Landscape Ecology 31(5), 1117–1137, mu doi: 10.1007/s10980-015-0323-0.

Nelken, I., Rotman, Y., and Yosef, O. B. (1999). “Responses of auditory-cortex neurons to structural features of natural sounds,” Nature 397(6715), 154–157, mu doi: 10.1038/16456.

Pan, W., Goodale, E., Jiang, A., and Mammides, C. (2024). “The effect of latitude on the efficacy of acoustic indices to predict biodiversity: A meta-analysis,” Ecological Indicators 159, 111747, mu doi: 10.1016/j.ecolind.2024.111747.

Park, S., Salles, A., Allen, K., Moss, C. F., and Elhilali, M. (2021). “Natural Statistics as Inference Principles of Auditory Tuning in Biological and Artificial Midbrain Networks,” eneuro 8(3), ENEURO.0525–20.2021, mu doi: 10.1523/ENEURO.0525-20.2021.

Phillips, Y. F., Towsey, M., and Roe, P. (2018). “Revealing the ecological content of long-duration audio-recordings of the environment through clustering and visualisation,” PLOS ONE 13(3), e0193345, mu doi: 10.1371/journal.pone.0193345.

Pijanowski, B. C., Villanueva-Rivera, L. J., Dumyahn, S. L., Farina, A., Krause, B. L., Napoletano, B. M., Gage, S. H., and Pieretti, N. (2011). “Soundscape Ecology: The Science of Sound in the Landscape,” BioScience 61(3), 203–216, mu doi: 10.1525/bio.2011.61.3.6.

Rees, A., and Malmierca, M. S. (2005). “Processing of Dynamic Spectral Properties of Sounds,” in International Review of Neurobiology, 70 (Elsevier), pp. 299–330, mu doi: 10.1016/S0074-7742(05)70009-X.

Richards, D. G., and Wiley, R. H. (1980). “Reverberations and Amplitude Fluctuations in the Propagation of Sound in a Forest: Implications for Animal Communication,” The American Naturalist 115(3), 381–399, mu doi: 10.1086/283568.

Rieke, F., Bodnar, D. A., and Bialek, W. (1995). “Naturalistic stimuli increase the rate and efficiency of information transmission by primary auditory afferents,” Proceedings of the Royal Society of London. Series B: Biological Sciences 262(1365), 259– 265, mu doi: 10.1098/rspb.1995.0204.

Rolland, J., and Freeman, B. G. (2023). The Latitudinal Diversity Gradient (Oxford University Press), mu doi: 10.1093/obo/9780199941728-0144.

Römer, H. (1998). “The Sensory Ecology of Acoustic Communication in Insects,” in Comparative Hearing: Insects, edited by R. R. Fay, A. N. Popper, R. R. Hoy, A. N. Popper, and R. R. Fay, 10 (Springer New York, New York, NY), pp. 63–96, mu doi: 10.1007/978-1-4612-0585-2_3.

Römer, H. (2001). “Ecological Constraints for Sound Communication: From Grasshoppers to Elephants,” in Ecology of Sensing, edited by F. G. Barth and A. Schmid (Springer Berlin Heidelberg, Berlin, Heidelberg), pp. 59–77, mu doi: 10.1007/978-3-662-22644-5_4.

Ross, S. R. P. J., Friedman, N. R., Dudley, K. L., Yoshimura, M., Yoshida, T., and Economo, E. P. (2018). “Listening to ecosystems: Data-rich acoustic monitoring through landscape-scale sensor networks,” Ecological Research 33(1), 135–147, mu doi: 10.1007/s11284-017-1509-5.

Sawilowsky, S. S. (1990). “Nonparametric Tests of Interaction in Experimental Design,” Review of Educational Research 60(1), 91–126, mu doi: 10.3102/00346543060001091.

Sawilowsky, S. S., Blair, R. C., and Higgins, J. J. (1989). “An Investigation of the Type I Error and Power Properties of the Rank Transform Procedure in Factorial ANOVA,” Journal of Educational Statistics 14(3), 255, mu doi: 10.2307/1165018.

Schreiner, C. E., and Urbas, J. V. (1986). “Representation of amplitude modulation in the auditory cortex of the cat. I. The anterior auditory field (AAF),” Hearing Research 21(3), 227–241, mu doi: 10.1016/0378-5955(86)90221-2.

Senter, P. (2008). “Voices of the past: A review of Paleozoic and Mesozoic animal sounds: Review,” Historical Biology 20(4), 255–287, mu doi: 10.1080/08912960903033327.

Shannon, R. V., Zeng, F.-G., Kamath, V., Wygonski, J., and Ekelid, M. (1995). “Speech Recognition with Primarily Temporal Cues,” Science 270(5234), 303–304, mu doi: 10.1126/science.270.5234.303.

Simoncelli, E. P., and Olshausen, B. A. (2001). “Natural Image Statistics and Neural Representation,” Annual Review of Neuroscience 24(1), 1193–1216, mu doi: 10.1146/annurev.neuro.24.1.1193.

Singh, N. C., and Theunissen, F. E. (2003). “Modulation spectra of natural sounds and ethological theories of auditory processing,” The Journal of the Acoustical Society of America 114(6), 3394– 3411, mu doi: 10.1121/1.1624067.

Skerritt-Davis, B., and Elhilali, M. (2021a). “Computational framework for investigating predictive processing in auditory perception,” Journal of Neuroscience Methods 360, 109177, mu doi: 10.1016/j.jneumeth.2021.109177.

Skerritt-Davis, B., and Elhilali, M. (2021b). “Neural Encoding of Auditory Statistics,” The Journal of Neuroscience 41(31), 6726– 6739, mu doi: 10.1523/JNEUROSCI.1887-20.2021.

Smith, E. C., and Lewicki, M. S. (2006). “Efficient auditory coding,” Nature 439(7079), 978–982, mu doi: 10.1038/nature04485.

Stebbins, W. C., and Moody, D. B. (1994). “How Monkeys Hear the World: Auditory Perception in Nonhuman Primates,” in Comparative Hearing: Mammals, edited by R. R. Fay, A. N. Popper, R. R. Fay, and A. N. Popper, 4 (Springer New York, NewYork, NY), pp. 97–133, mu doi: 10.1007/978-1-4612-2700-7_4.

Stein, R. C. (1968). “Modulation in Bird Sounds,” The Auk 85(2), 229–243, mu doi: 10.2307/4083583.

Sueur, J., and Aubin, T. (2003). “Specificity of cicada calling songs in the genus Tibicina (Hemiptera: Cicadidae),” Systematic Entomology 28(4), 481–492, mu doi: 10.1046/j.1365-3113.2003.00222.x.

Sueur, J., and Farina, A. (2015). “Ecoacoustics: The Ecological Investigation and Interpretation of Environmental Sound,” Biosemiotics 8(3), 493–502, mu doi: 10.1007/s12304-015-9248-x.

Sueur, J., Pavoine, S., Hamerlynck, O., and Duvail, S. (2008). “Rapid Acoustic Survey for Biodiversity Appraisal,” PLoS ONE 3(12), e4065, mu doi: 10.1371/journal.pone.0004065.

Theunissen, F. E. (2018). “Biosound. a python library to analyze sounds used in communication” muhttps://hdl.handle.net/21.11116/0000-0004-4E23-E.

Theunissen, F. E., and Elie, J. E. (2014). “Neural processing of natural sounds,” Nature Reviews Neuroscience 15(6), 355–366, mu doi: 10.1038/nrn3731.

Thoret, E., Varnet, L., Boubenec, Y., Férriere, R., Le Tourneau, F.-M., Krause, B., and Lorenzi, C. (2020). “Characterizing amplitude and frequency modulation cues in natural soundscapes: A pilot study on four habitats of a biosphere reserve,” The Journal of the Acoustical Society of America 147(5), 3260–3274, mu doi: 10.1121/10.0001174.

Tishechkin, D. Yu. (2013). “Vibrational background noise in herbaceous plants and its impact on acoustic communication of small Auchenorrhyncha and Psyllinea (Homoptera),” Entomological Review 93(5), 548–558, mu doi: 10.1134/S0013873813050035.

Tolhurst, D. J., Tadmor, Y., and Chao, T. (1992). “Amplitude spectra of natural images,” Ophthalmic and Physiological Optics 12(2), 229–232, mu doi: 10.1111/j.1475-1313.1992.tb00296.x.

Torralba, A., and Oliva, A. (2003). “Statistics of natural image categories,” Network: Computation in Neural Systems 14(3), 391–412, mu doi: 10.1088/0954-898X_14_3_302.

Tsandilas, T., and Casiez, G. (2025). “The illusory promise of the aligned rank transform,” [preprint] https://statransform.github.io/jov.

Varnet, L., Ortiz-Barajas, M. C., Erra, R. G., Gervain, J., and Lorenzi, C. (2017). “A cross-linguistic study of speech modulation spectra,” The Journal of the Acoustical Society of America 142(4), 1976–1989, mu doi: 10.1121/1.5006179.

Venezia, J. H., Hickok, G., and Richards, V. M. (2016). “Auditory “bubbles”: Efficient classification of the spectrotemporal modulations essential for speech intelligibility,” The Journal of the Acoustical Society of America 140(2), 1072–1088, mu doi: 10.1121/1.4960544.

Viemeister, N. F. (1979). “Temporal modulation transfer functions based upon modulation thresholds,” The Journal of the Acoustical Society of America 66(5), 1364–1380, mu doi: 10.1121/1.383531.

Voss, R. F., and Clarke, J. (1975). “‘1/f noise’ in music and speech,” Nature 258(5533), 317–318, mu doi: 10.1038/258317a0.

Wallaert, N., Varnet, L., Moore, B. C. J., and Lorenzi, C. (2018). “Sensorineural hearing loss impairs sensitivity but spares temporal integration for detection of frequency modulation,” The Journal of the Acoustical Society of America 144(2), 720–733, mu doi: 10.1121/1.5049364.

Waser, P. M., and Brown, C. H. (1986). “Habitat acoustics and primate communication,” American Journal of Primatology 10(2), 135–154, mu doi: 10.1002/ajp.1350100205.

Wiley, R. H., and Richards, D. G. (1978). “Physical constraints on acoustic communication in the atmosphere: Implications for the evolution of animal vocalizations,” Behavioral Ecology and Sociobiology 3(1), 69–94, mu doi: 10.1007/BF00300047.

Wobbrock, J. O., Findlater, L., Gergle, D., and Higgins, J. J. (2011). “The aligned rank transform for nonparametric factorial analyses using only anova procedures,” in Proceedings of the SIGCHI Conference on Human Factors in Computing Systems, ACM, Vancouver BC Canada, pp. 143–146, mu doi: 10.1145/1978942.1978963.

Woolley, S. M. N., Fremouw, T. E., Hsu, A., and Theunissen, F. E. (2005). “Tuning for spectro-temporal modulations as a mechanism for auditory discrimination of natural sounds,” Nature Neuroscience 8(10), 1371–1379, mu doi: 10.1038/nn1536.

Yang, M., De Coensel, B., and Kang, J. (2015). “Presence of 1/f noise in the temporal structure of psychoacoustic parameters of natural and urban sounds,” The Journal of the Acoustical Society of America 138(2), 916–927, mu doi: 10.1121/1.4927033

Zina, J., and Haddad, C. F. B. (2005). “Reproductive activity and vocalizations of Leptodactylus labyrinthicus (Anura: Leptodactylidae) in southeastern Brazil,” Biota Neotropica 5(2), 119– 129, mu doi: 10.1590/S1676-06032005000300008.

